# Molecular Signatures and Cellular Diversity During Mouse Habenula Development

**DOI:** 10.1101/2022.02.21.481251

**Authors:** Lieke L. van de Haar, Danai Riga, Juliska E. Boer, Youri Adolfs, Thomas E. Sieburgh, Roland E. van Dijk, Kyoko Watanabe, Nicky C.H. van Kronenburg, Mark H. Broekhoven, Danielle Posthuma, Frank J. Meye, Onur Basak, R. Jeroen Pasterkamp

**Affiliations:** Department of Translational Neuroscience, UMC Utrecht Brain Center, University Medical Center, Utrecht University, Utrecht, 3584 CG, The Netherlands; Department of Complex Trait Genetics, Center for Neurogenomics and Cognitive Research, Neuroscience Campus Amsterdam, VU University Amsterdam, Amsterdam, 1081 HV, The Netherlands

**Keywords:** Habenula, mouse, spatial expression, development, single-cell RNA sequencing, electrophysiology, neuronal subtype, axon guidance, neurotransmitter, connectivity

## Abstract

The habenula plays a key role in various motivated and pathological behaviors and is composed of molecularly distinct neuron subtypes. Despite progress in identifying mature habenula neuron subtypes, how these subtypes develop and organize into functional brain circuits remains largely unknown. Here we performed single-cell transcriptional profiling of mouse habenular neurons at critical developmental stages instructed by detailed three-dimensional anatomical data. Our data reveal cellular and molecular trajectories during embryonic and postnatal development leading to different habenular subtypes. Further, based on this analysis our work establishes the distinctive functional properties and projection target of a previously uncharacterized subtype of *Cartpt^+^* habenula neurons. Finally, we show how comparison of single-cell transcriptional profiles and GWAS data links specific developing habenular subtypes to psychiatric disease. Together, our study begins to dissect the mechanisms underlying habenula neuron subtype-specific development and creates a framework for further interrogation of habenular development in normal and disease states.

## INTRODUCTION

The habenula (Hb) is an evolutionary conserved brain nucleus that plays a key role in processing reward information and mediating aversive responses to negative stimuli. It receives input from limbic areas and the basal ganglia through the stria medularis (SM), while its efferent projections form the fasciculus retroflexus (FR) and connect to mid- and hindbrain regions. The Hb has been implicated in several human diseases, such as major depressive disorder (MDD) and schizophrenia (for review see (Hu et al., 2020)). How such a relatively small group of neurons underpins a disproportionally large number of (patho)physiological functions is incompletely understood but may be explained by their heterogeneous nature. Historically, the Hb is subdivided into two subnuclei based on anatomical features and reported functions: the medial Hb (MHb) and the lateral Hb (LHb). The MHb contains substance P-ergic (SP-ergic) and cholinergic neurons, projects to the interpeduncular nucleus (IPN) and is linked to addiction, anxiety and depression (Aizawa et al., 2012; Fowler et al., 2011; Herkenham and Nauta, 1977; Molas et al., 2017; Yamaguchi et al., 2013; Zhang et al., 2016). The LHb contains predominantly glutamatergic neurons and projects to the rostromedial tegmental nucleus (RMTg), raphe nuclei, and ventral tegmental area (VTA) (Herkenham and Nauta, 1977; Quina et al., 2015). It coordinates responses to aversive stimuli, reward-processing, feeding and is implicated in addiction (e.g. (Flanigan et al., 2020; Hong et al., 2011; Lammel et al., 2012; Matsumoto and Hikosaka, 2007; Meye et al., 2016; Proulx et al., 2014; Tian and Uchida, 2015)). Several studies have revealed further subdivisions of the Hb by identifying clusters of Hb neurons with select molecular signatures. Recent single-cell transcriptional profiling studies have defined at least nine molecularly distinct subtypes of neurons in the adult mouse Hb, while also the adult zebrafish Hb was reported to contain multiple transcriptionally distinct neuron populations (Aizawa et al., 2012; Hashikawa et al., 2020; Pandey et al., 2018; Wagner et al., 2016; Wallace et al., 2020). Interestingly, some of these molecularly defined subtypes are specifically engaged in foot shock responses (Hashikawa et al., 2020) or linked to specific neural activity (Pandey et al., 2018), denoting the functional implications of their diversity. Despite this progress, the functional properties and connectivity patterns of most molecularly distinct subtypes of mammalian Hb neurons remain unknown.

Mouse Hb neurons originate from progenitors in prosomere 2 (P2) between E10.5 and E18.5 (Guo and Li, 2019; Vue et al., 2007; Wong et al., 2018). After mitosis, newly born neurons leave the ventricular zone (VZ) and migrate laterally into the presumptive Hb (Ruiz-reig et al., 2019). Upon reaching their final location, Hb neurons send out axons instructed by molecular cues in the environment (Funato et al., 2000; Ichijo and Toyama, 2015; Schmidt et al., 2014). However, while the functional roles and connectivity patterns of the adult mammalian MHb and LHb have been studied in great detail, much less is known about the mechanisms underlying mouse Hb morphogenesis (for review see (Schmidt and Pasterkamp, 2017)). Further, how distinct subtypes of MHb and LHb neurons emerge and develop, or contribute to disease remains largely unexplored.

Here, we combine single-cell transcriptional profiling of the developing mouse Hb with molecular, anatomical and functional approaches to unveil developmental trajectories leading to different subtypes of adult Hb neurons. Using these data, we study the developmental gene expression patterns of candidate gene families, identify and characterize a subtype of poorly defined *Cartpt^+^* Hb neurons, and link clusters of developing Hb neurons to genetic risk variants of psychiatric disease. Together, our data constitute a framework for future studies on Hb development in health and disease.

## RESULTS

### Three-dimensional analysis of habenula development

To study the development of neuronal heterogeneity in the mouse Hb we aimed to perform single-cell RNA sequencing (scRNAseq) at key developmental timepoints. To supplement published, but fragmented, data on Hb development, we performed whole brain immunohistochemistry, 3D imaging of solvent-cleared organ (3DISCO) tissue clearing, and fluorescent light sheet microscopy (FLSM) (Adolfs et al., 2021). These data were then used to aid selection of developmental stages for scRNAseq (Fig. 1A, S1).

**Figure 1.**
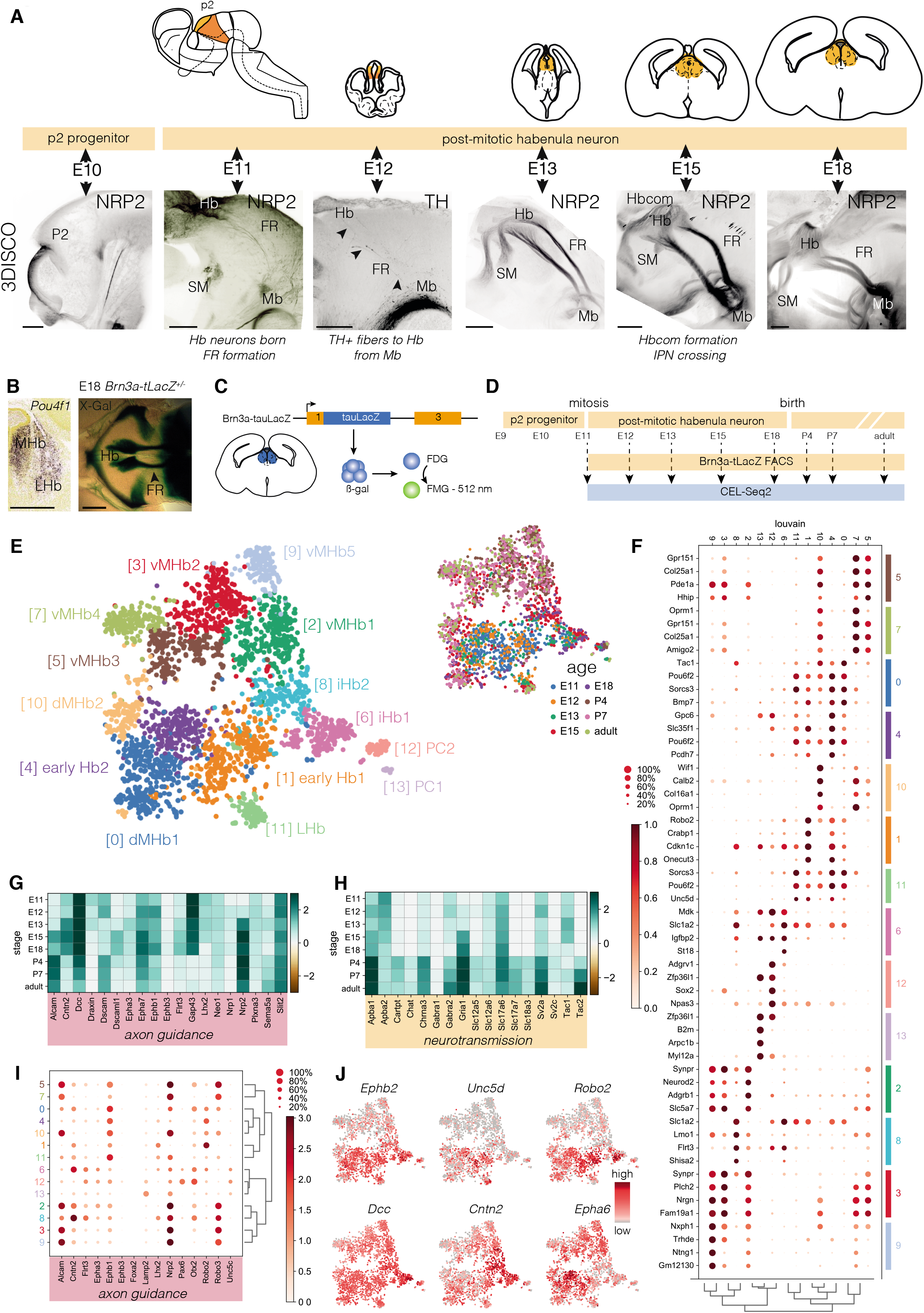
Analysis of the developing mouse habenula using FLSM and scRNAseq. **A,** Top: Schematic representation of the developing mouse Hb. Bottom: Whole-mount immunostaining for NRP2 or TH of mouse embryos followed by 3DISCO and FLSM to visualize the developing Hb and its connections. Caudal is to the right, dorsal to the top. **B,** Left: *In situ* hybridization for *Pou4f1/Brn3a* in the E18 mouse Hb (from Allen Brain Atlas). Right: Wholemount β-gal staining. Top view. **C,** Strategy for labeling *Brn3a-tLacZ^+^* neurons for FACS by β-gal-mediated hydrolysis of FDG to fluorescent FMG. **D,** Developmental and experimental timelines showing sampling stages for CEL-Seq2. **E,** t-SNE embedding showing clusters as detected by the Louvain algorithm per single cell. Right: t-SNE embedding color-coded by developmental stage. d, dorsal; i, immature; PC, progenitor cell; v, ventral. **F,** Dot-plot of the scaled and normalized expression of the top 4 DEG per cluster calculated with Wilcoxon rank sum test after filtering (minimum normalized expression within cluster = 0.25, maximum normalized expression in other groups = 0.5, minimum fold change = 2). **G, H,** Matrix plot of selected axon guidance- and neurotransmission-related genes per developmental stage. **I**, Dot-plot of the normalized expression of axon guidance-related genes per cluster. **J**, t-SNE embedding showing the expression axon guidance-related genes. Scale bar: 200 μm (A), 400 μm (B). See Figure S1-5.

Neuropilin-2 (NRP2), roundabout-1 (ROBO1) and ROBO3 were used to label Hb neurons and projections (Belle et al., 2014; Giger et al., 2000). Tyrosine hydroxylase (TH) enabled visualization of the midbrain (Mb) dopamine system, an important Hb target and source of afferent projections (Schmidt et al., 2014). From E11 onwards NRP2^+^ neurons were detected in P2. These Hb neurons sent out NRP2^+^/ROBO1^+^/3^+^ axons towards the midbrain forming the FR. At E12, FR axon number had increased and TH^+^ pioneer axons extended to the LHb (Schmidt et al., 2014). At E13, the NRP2^+^ SM projected into the Hb and the first NRP2^+^/ROBO3^+^ axons crossed the midline of the Mb (Fig. S1). At E15, the number of crossing axons had increased, TH^+^ axons were restricted to the FR sheath region, and the NRP2^+^/ROBO1^+^ Hb commissure (Hbcom) formed. At E18, Hb size and connectivity had further increased. Based on these data and available literature, different embryonic stages (E11-E13, E15, E18) were selected for scRNAseq. P4 and P7 were included to cover processes such as synaptogenesis, and an adult stage to enable trajectory analysis towards adult subtypes.

### scRNAseq identifies different subtypes of habenula neurons during development

To specifically isolate Hb neurons, fluorescence activated cell sorting (FACS) of LacZ^+^ neurons from *Brn3a-tauLacZ* mice was performed (Fig. 1B-D, S2A, B). *Brn3a*, also known as *Pou4f1*, is a marker of post-mitotic Hb neurons (Raisa Eng et al., 2001). FACS-based CEL-seq2 scRNAseq resulted in transcriptional profiles of 5,756 cells across eight ages expressing 38,068 genes (Fig. 1D, S2C-D, S3A-C). Louvain clustering resulted in 14 distinct clusters containing cells derived from different developmental stages (Fig. 1E, S3D-F). Based on the expression of published marker genes (Chatterjee et al., 2015; Hashikawa et al., 2020; Lee et al., 2017; Lipiec et al., 2019; Wagner et al., 2016; Wallace et al., 2020), we annotated *Vim^+^/Sox2^+^* progenitors (PC; cluster 12 and 13), *Dcc^+^/Cntn2^+^* immature Hb (iHb) neurons (6 and 8), *Gap43^+^/Un5d^+^* LHb neurons (11), *Chat^+^* cholinergic ventral MHb neurons (2, 3, 5, 7 and 9) and *Tac1^+^* SP-ergic dorsal MHb neurons (0 and 10) (Fig. 1E, F, Table S1). The remaining *Robo2^+^* (1) and *Pcdh7^+^* (1 and 4) clusters predominantly contained E11 and E12 neurons. To examine which cellular subtypes were present per developmental stage, we performed Louvain clustering per stage (Fig. S4). This revealed between 5-10 subtypes per stage, each with cluster-specific gene expression patterns. Together, these results unveil a high level of cellular heterogeneity in the developing mouse Hb.

The generated Hb scRNAseq dataset (DevHb) enables in depth analysis of the temporal and subtype-specific expression of candidate genes that instruct important cellular processes. To highlight this potential, we examined the temporal expression of axon guidance and neurotransmission genes crucial for Hb development and function (Fig. 1G, H, S5A). This revealed, for example, that expression of the axon guidance receptor *Dcc* decreased towards adulthood, while *Nrp2* and *Alcam* expression increased. Further, members of the same axon guidance receptor families often displayed very distinct temporal expression patterns (e.g. EPH receptors). Most neurotransmission-related transcripts were expressed from postnatal stages onwards, although some genes were already detected at E11 (e.g. *Tac1, Scl17a6*)(Fig. 1H, S5A). In addition, several of the selected genes displayed cluster-specific or -enriched expression throughout development (e.g. *Tac1, Ephb2*) (Fig. 1F, I, J, S4A, S5B). In all, this analysis revealed unique spatiotemporal and subtype-specific expression patterns that will help to explain how Hb neuron subtypes develop and mature.

### Cellular heterogeneity is established during postnatal development

Next, we determined how the DevHb clusters relate to adult Hb clusters reported previously (Hashikawa et al., 2020;Wallace et al., 2020). First, the two adult datasets were compared and a strong correlation was found (r = 0.8-0.95) (Fig. S6A). Correlation analysis of the DevHb data to the adult datasets showed that DevHb MHb populations correlated to the reported adult MHb clusters, while DevHb LHb cluster 11 correlated to all adult LHb populations (Fig. 2A-C). As expected, DevHb clusters containing PCs or iHb (6, 8, 12 and 13) did not correlate well with adult clusters. Using these results and expression of previously determined cluster-specific marker genes (Hashikawa et al., 2020; Wagner et al., 2016; Wallace et al., 2020), we next determined for each dataset the cluster(s) that contained a specific Hb neuron subtype. This showed that the DevHb dataset captured the previously reported molecular heterogeneity of the adult MHb. In contrast, only one LHb cluster was found in the DevHb, while multiple adult LHb have been reported (Fig. 2D). This is presumably a result of using the *Brn3a-tauLacZ* mouse in which the MHb is more extensively labeled then the LHb (Fig. S2B).

**Figure 2.**
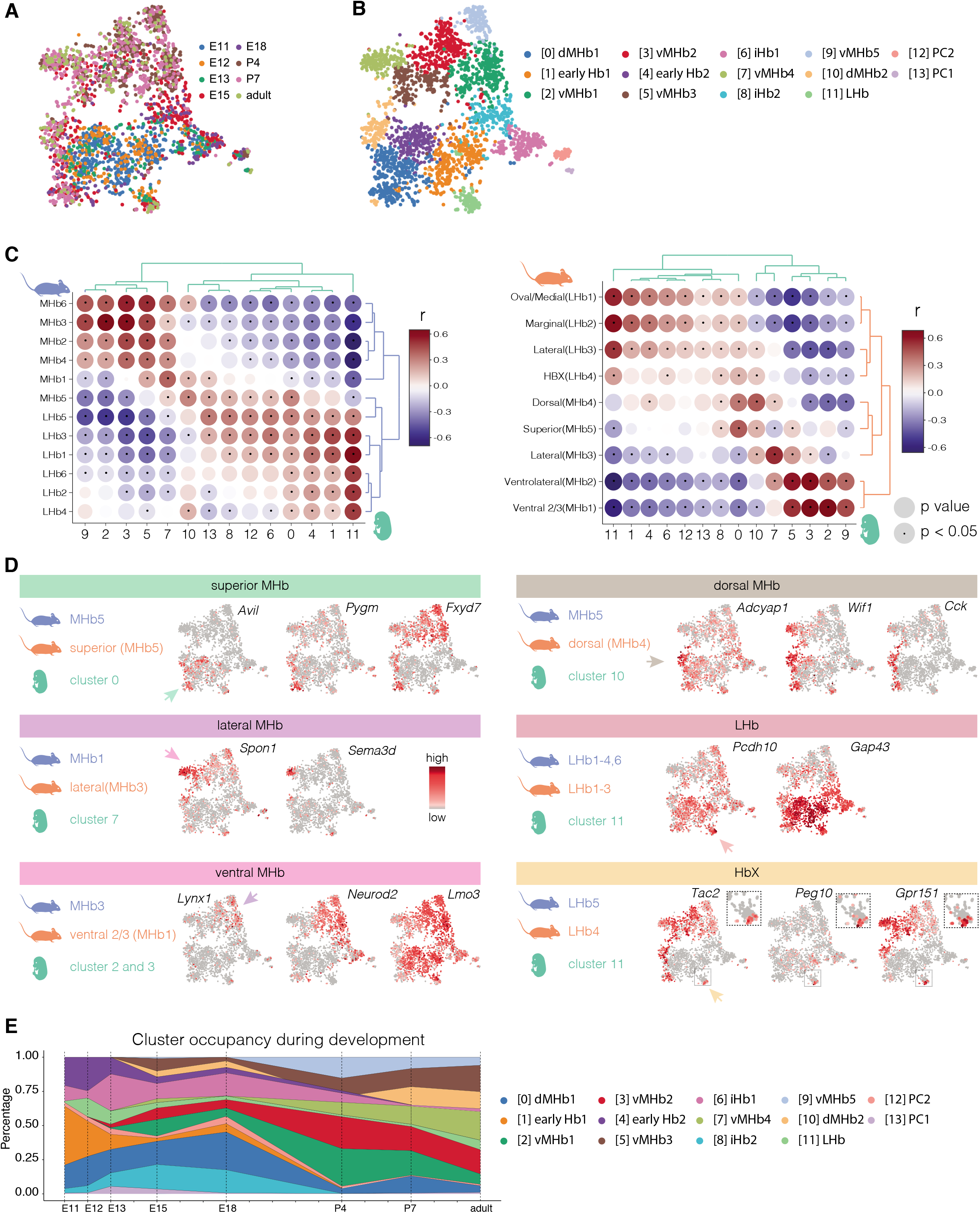
Neuronal heterogeneity in the mouse habenula is established at early postnatal stages. **A, B,** Color-coding of t-SNE embedding of the DevHb data by developmental stage (A) and Louvain cluster (B). **C,** Correlation matrices showing the correlation (r) between the clusters of the DevHb dataset (green) and the Hashikawa (blue) or Wallace dataset (orange). Disc size indicates *P* value and a dot statistical significance (*P* < 0.05). **D,** Overview of Hb cell clusters indicated with marker gene expression levels on t-SNE. For each dataset the cluster is indicated that contains the specific Hb subtype (blue: Hashikawa, orange: Wallace, green: DevHb). Arrow indicates DevHb cluster. **E,** Occupancy plot shows relative abundance of Louvain clusters per developmental stage in the DevHb dataset. See Figure S6.

No previous scRNAseq data were available on the developing mouse Hb, but scRNAseq was recently performed at one zebrafish larval stage (Pandey et al., 2018). Correlation of the DevHb clusters to larval and adult zebrafish data showed that mouse MHb clusters correlate to dorsal Hb zebrafish clusters and the mouse LHb cluster to ventral Hb zebrafish clusters (Fig. S6B-D). Further, iHb populations (6 and 8) strongly correlated to the immature larval zebrafish Hb population (La_Hb13), while mouse PCs (12 and 13) correlated to adult zebrafish PCs (Ad_16) (Fig. S6C, D). Interestingly, the heterogeneity found in the adult zebrafish Hb was already present at 10 dpf, a stage analogous to mouse postnatal stages (Fig. S6B). To determine when Hb heterogeneity is established, cluster occupancy in the DevHb data was plotted per developmental stage (Fig. 2E). PC and iHb populations were primarily abundant between E11 and E18, and mostly absent by P4. Clusters that strongly correlated to adult clusters appeared latest at P4 (0-3, 5, 7, 9-11). Correlation of subclusters per stage (Fig. S4) showed that adult mouse clusters correlated most strongly to postnatal and not embryonic subclusters (Fig. S6E). Together these data for the first time establish that the heterogeneity of the adult mouse Hb is established at early postnatal stages.

### Developmental trajectories of mouse habenula neurons are marked by specific gene expression patterns

Next, we wanted to identify the cellular and molecular programs that instruct Hb subtype development. Analysis of the DevHb data using CellRank trajectory inference identified PC cluster 13 as the root of the Hb developmental trajectory (Fig. 3A-C). From PC13, a decrease of the early marker genes *Id3* and *Vim* was found along latent time followed by a trajectory to iHb (yellow) (cluster 6), marked by increased *Cntn2* and *Dcc* expression. Then the trajectory split, leading to LHb neurons, and dorsal and ventral subsets of MHb neurons. Dorsal MHb subsets were defined by *Calb2*. The dorsal MHb-superior (dMHb-S) trajectory led to *Tac1^+^* SP-ergic neurons, while dorsal MHb-dorsal (dMHb-D) neurons were *Calb1^+^* (Fig. 3A, B). Both ventral MHb trajectories were marked by the cholinergic marker *Chat*, but a distinction could be made between *Spon1^+^* ventral MHb-lateral (vMHb-L) and *Synpr^+^* ventral MHb-ventral (vMHb-V) neurons (Fig. 3A-C). These CellRank-determined trajectory endpoints matched the final cell types defined by correlation of the DevHb to adult clusters (Fig. 2D, E). Interestingly, according to CellRank latent time, cluster occupancy analysis and ISH, *Tac1^+^* SP-neurons (cluster 0, MHb-S, E11) arise before *Chat^+^* cholinergic neurons (cluster 9, vMHb, E18)(Fig. 2E, 3D, E).

**Figure 3.**
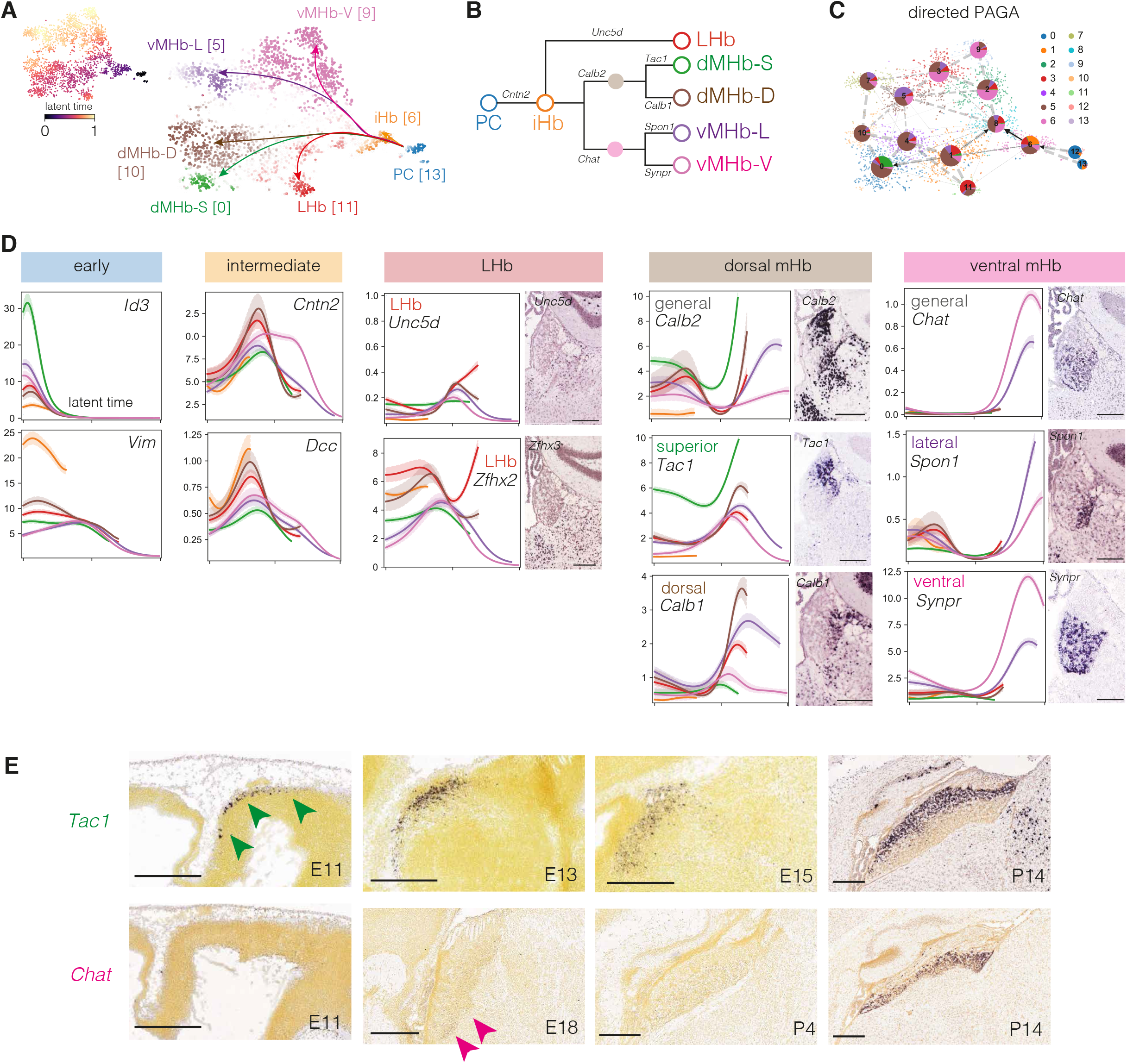
Trajectory inference reveals multiple developmental trajectories. **A, B**, CellRank trajectory inference analysis of the DevHb data identifies one initial (PC) and one intermediate state (iHb), and five trajectory end-points, plotted on t-SNE embedding. [n] Louvain cluster (A) or represented as dendrogram based (including marker genes) (B). **C,** PAGA showing the likelihood per cell within a cluster to reach a terminal state (i.e. trajectory endpoint). Arrows indicate direction of differentiation. **D,** Marker gene expression trend per trajectory position along latent time color-coded by trajectory. Adult coronal ISH images from Allen Brain Atlas (ABA). **E,** ISH on sagittal sections. Green arrowheads, *Tac1* in superior medial Hb. Pink arrows, *Chat* in ventral medial Hb (ABA). Scale bar, 200 μm (D), 400 μm (E).

To identify molecular signatures of the maturational trajectories and endpoints, expression trends for the top 50 most highly associated genes per trajectory were plotted along latent time (Fig. 4A, B). Known PC genes (*Dbi*, *Id3* and *Vim*) decreased on the trajectory from PC to iHb. Additionally, several trajectory-specific transcripts peaked along latent time, e.g. *Dab1* (to LHb), *Tenm3* and *Pou6f2* (to dMHb-D), *March1* (to dMHb-S), *Nkain3* (to vMHb-V) and *Ebf1* (to vMHb-L). Transcripts peaking at trajectory endpoints were mostly trajectory-specific but in some cases overlapping (e.g. *Syt6* and *Kcng4* marked both dMHb endpoints). We also assessed a possible role of specific “master” transcription factors (TFs) and effector genes (together defined as a regulon). Regulon activity was primarily associated with developmental stage but not cellular subtype, with the exception of PCs (Fig. 4C-E, S7). Interestingly, several regulons displayed specific temporal expression, e.g. early regulons (*Onecut3*, *Cux2*) and late regulons (*Mef2d*) with reported roles in embryonic development and synaptic plasticity, respectively (Fig. 4D, E). Together, these analyses identify genes that mark, and may instruct, developmental trajectories to adult neuronal subtypes and unveil temporally expressed regulons in Hb neuron development and maturation.

**Figure 4.**
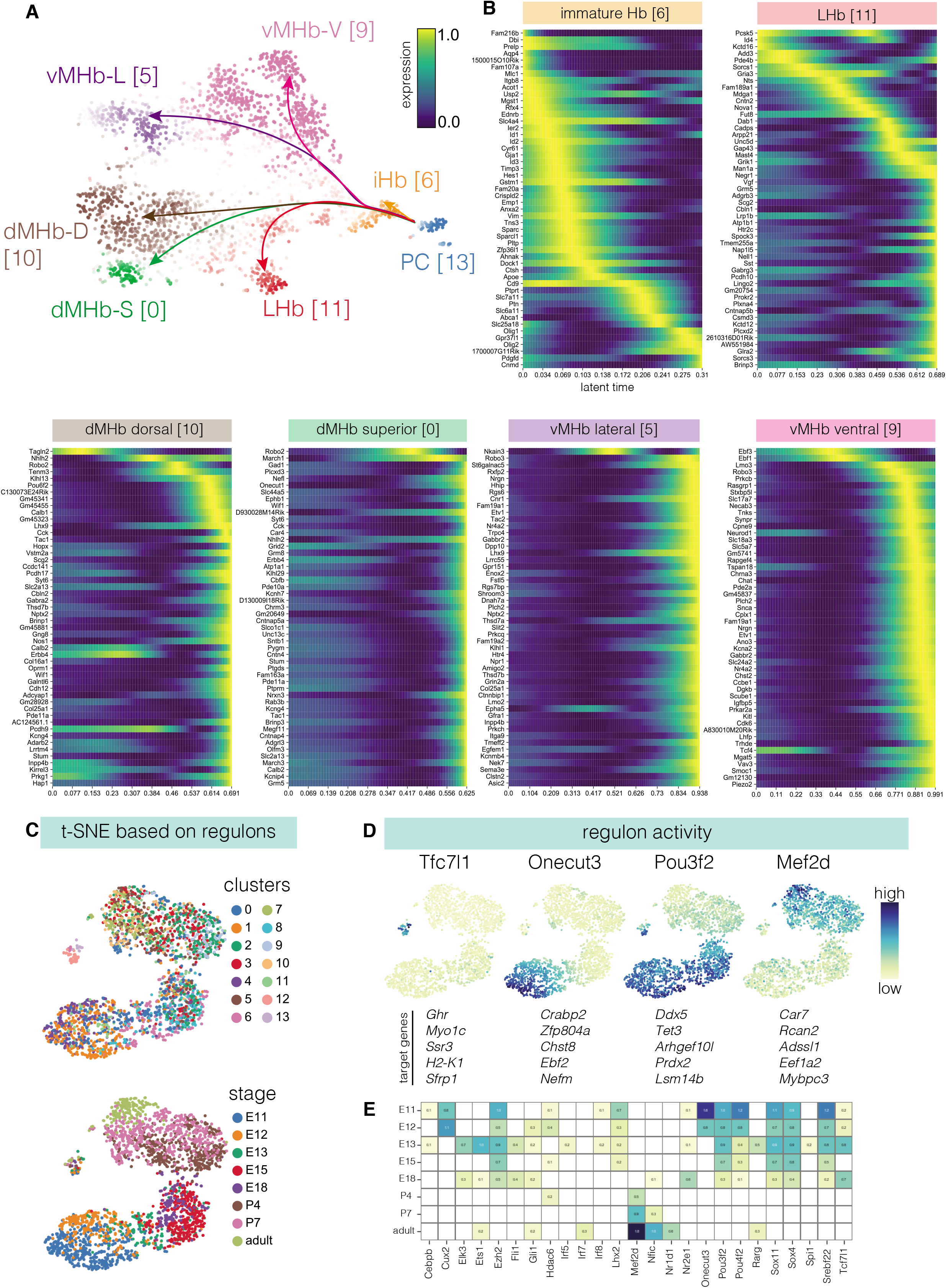
Gene expression signatures underlying habenula neuron subtype development. **A,** CellRank trajectory inference analysis DevHb data (as in Fig. 3A). **B,** Heatmaps showing temporal expression patterns (calculated using a general additive model) of the top 50 most highly associated genes per trajectory sorted according to their peak at latent time. **C,** t-SNE embedding based on regulon expression computed with pySCENIC, color-coded by Louvain cluster (top) or developmental stage (bottom). **D,** Regulon expression plotted on novel t-SNE embedding (top) including target genes per regulon (bottom). **E,** Heatmap of z-scores of detected regulons per timepoint. See Figure S7.

### Dissection of early habenula (non)neuronal development

The DevHb dataset primarily contains post-mitotic Hb neurons because of the use of *Brn3a-tauLacZ* mice. To examine in more in detail (early) Hb development and non-neuronal subtypes, scRNAseq was performed on the whole E18 Hb. Cluster occupancy analysis indicated that at E18 cells with different maturational stages are present (Fig. 2E).

In many species the Hb exhibits left-right asymmetry. We therefore collected left and right Hb and determined DEGs. In total 948 cells were analyzed that expressed a total of 28,237 genes. 5 DEGs were found in the right Hb and 11 DEGs in the left Hb (adjusted *P*-value < 0.05)(Fig. 5A, Table S2, S3). However, this differential gene expression was not confirmed by ISH (Allen Brain Atlas) and cells from the left and right Hb showed overlapping UMAP placement (Fig. 5B). As these data do not support strong asymmetry at the transcriptional level, no distinction was made between the left and right Hb in subsequent analyses (referred to as E18 WT). Louvain clustering identified nine distinct cell populations in the E18 WT data, which were annotated based on marker gene expression and localization (Fig. 5C, S8A-D). A large population of PCs was detected, which marked the trajectory starting point selected by scVELO (Fig. 5D), as observed for the DevHb data (Fig. 3A, B). However, in contrast to the DevHb data, analysis of E18 WT data also showed trajectories to oligodendrocyte precursor cells (OPC), thalamus neurons (Th) (3), and immature (thalamic) neurons (iNeur) (6). Correlation analysis confirmed this difference, as no strong correlation of E18 WT clusters 3, 6 and 8 was found with DevHb clusters (Fig. 5E). Thus, the E18 WT dataset provides additional cellular insight into early Hb development and the generation of other (non)neuronal cell types in and around the Hb.

**Figure 5.**
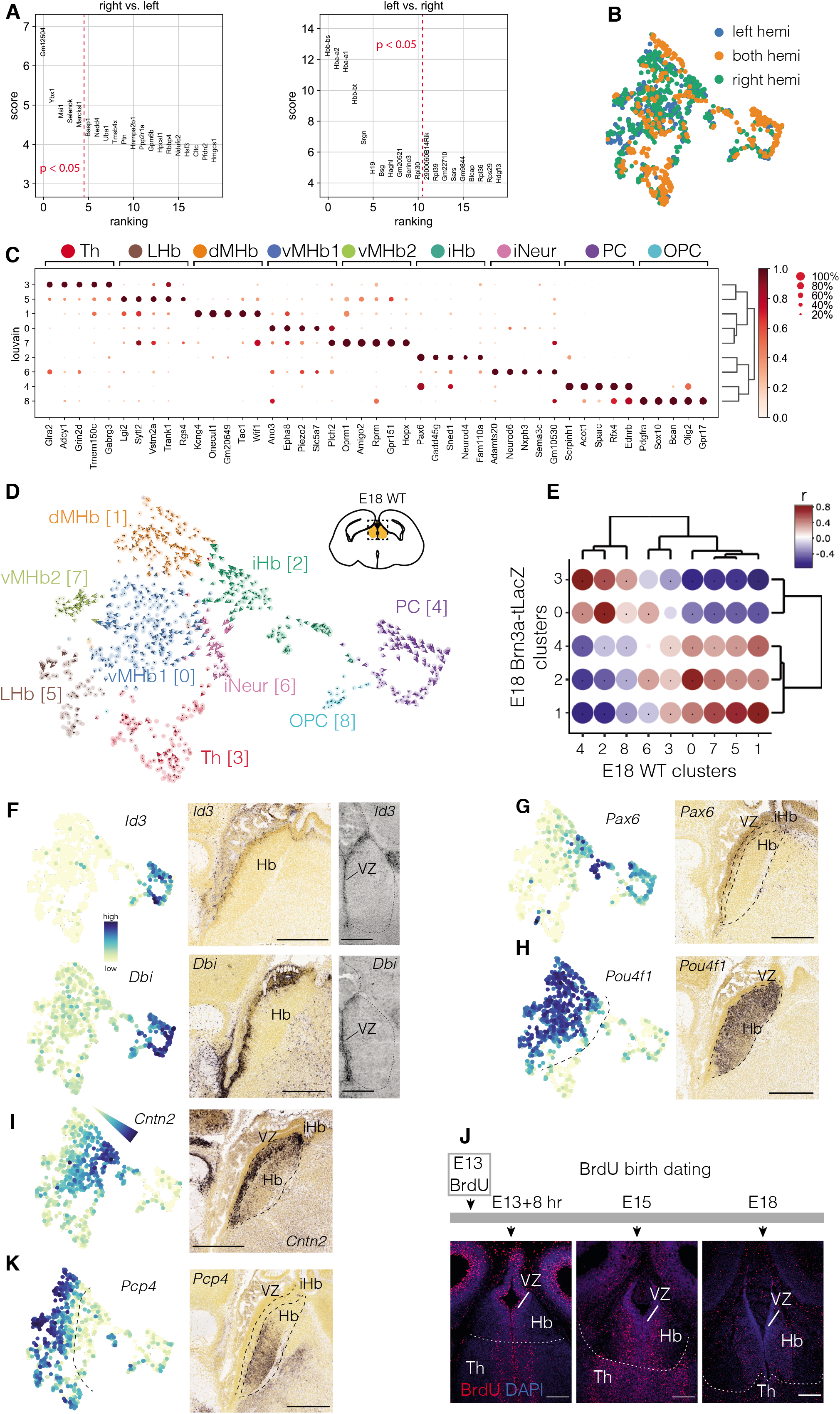
scRNAseq analysis of the whole E18 habenula. **A,** DEG between left and right Hb neurons calculated with the Wilcoxon Rank Sum test. Red dotted line, statistical significance threshold of adjusted *P* value < 0.05. **B,** Origin of cells (from left or right hemisphere) on UMAP. “Both hemi” indicates cells originating from a sample containing both hemispheres. **C,** Dot plot of the scaled and normalized expression of the top 5 DEG per cluster calculated with Wilcoxon rank sum test after filtering (minimum normalized expression within cluster = 0.5, maximum expression in other groups = 0.3, minimum fold change = 1.5). **D,** scVELO trajectory interference analysis of E18 mouse Hb cells (E18 WT). **E,** Correlation analysis of E18 WT and E18 Brn3a-tLacZ clusters (see Fig. S4). Disc size indicates *P* value and a dot statistical significance (*P* < 0.05). **(F, G, H, I, K)** Left: expression levels of indicated genes plotted on t-SNE embedding of E18 WT data. Right: ISH on sagittal or coronal sections of the E18 Hb (Allen Brain Atlas or this study). **(J)** BrdU was injected at E13, embryos were harvested at E13+8 hours, E15 and E18 and subjected to BrdU immunohistochemistry. Scale bar, 200 μm (F (coronal), J), 400 μm (F, G, H, I, K). See Figure S8.

Early development of the mouse Hb remains incompletely understood (Schmidt and Pasterkamp, 2017). Therefore, we analyzed the spatial expression of several genes that marked different parts of the trajectory from PCs to mature Hb neurons. ISH localized the PC markers *Id3* and *Dbi* to the VZ of the Hb lining the third ventricle (Fig. 5F, S8E). The Hb-specific trajectory leads from PCs into the iHb cluster (Fig. 5D). *Pax6*, essential for epithalamus development (Chatterjee et al., 2014), was expressed in PCs and a narrow adjacent region (iHb) (Fig. 5G). The iHb cluster was characterized by *Cntn2*, *Dcc*, *Flrt3* and *Pou4f1/Brn3a* (Fig. 5H, I, S8F). Trajectory inference analysis showed decreasing *Cntn2* expression along the trajectory towards more mature Hb clusters (Fig. 3D), which was confirmed by ISH (Fig. 5I). *In vivo* BrdU pulse labeling confirmed that *Cntn2^high^* neurons adjacent to the VZ are young neurons and that older *Cntn2^low^* neurons are located more laterally (Fig. 5J). More lateral neurons expressed *Pcp4* (Fig. 5K). Thus, the transition from PCs to adult Hb neurons is a topographical process that is accompanied by sequential gene expression changes.

### *Cartpt* expression marks a subset of physiologically distinct habenula neurons

Our data and work of others identify molecularly distinct Hb subtypes. However, whether these transcriptional identities associate with specific connectivity patterns and functional properties remains largely unexplored. To address this point, we studied the spatial distribution of a large set of genes that label one or a few clusters (including neurotransmission-related genes (Fig. S5)). This analysis highlighted a specific population of *Cartpt^+^* neurons at the boundary of the MHb and LHb, known as HbX and border zone regions (Fig. 6A) (Wagner et al., 2016; Wallace et al., 2020). *Cartpt* encodes CART peptide which, for example, affects locomotor activity and has anti-depressant effects (Singh et al., 2021). Despite these potent effects, the developmental characteristics, projection targets and functional properties of *Cartpt^+^* Hb neurons are unknown.

**Figure 6.**
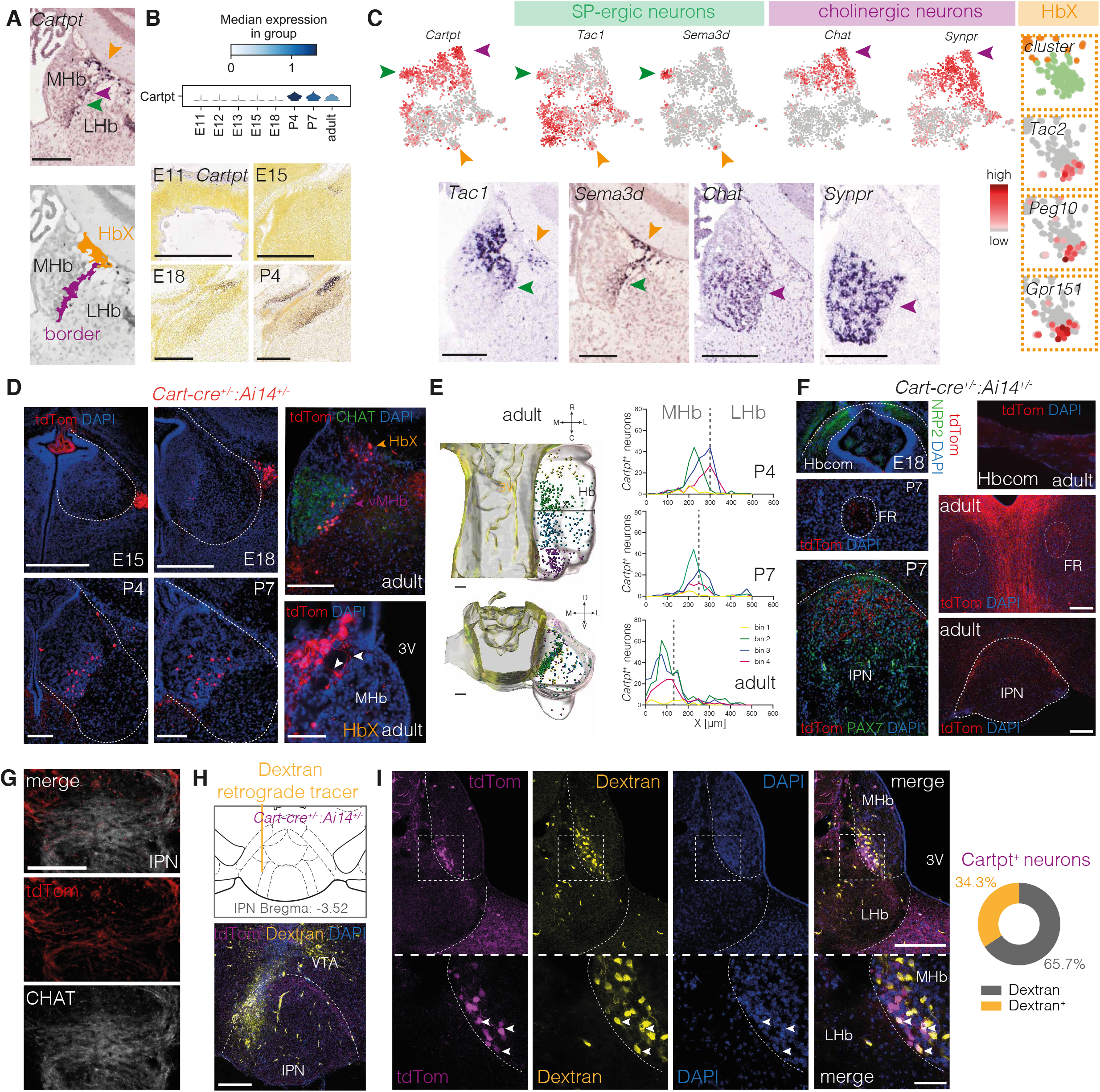
Molecular signature and projection target of Cartpt^+^ habenula neurons. **A,** Top: ISH for *Cartpt* in an E18 coronal section (Allen Brain Atlas (ABA)). Bottom: Location of the border and HbX regions. Arrowheads, different populations of neurons (see C). yellow, HbX; green, substance-P border; purple, cholinergic border. **B,** Top: violin plot of median *Cartpt* expression level per developmental stage in DevHb dataset. Bottom: ISH for *Cartpt* in sagittal sections (ABA). **C,** Top: gene expression plotted on t-SNE embedding. Bottom: ISH for indicated genes in adult coronal sections. **D,** Immunohistochemistry for tdTomato (and CHAT to mark vMHb) on coronal sections of *Cart-cre:Ai14* mice. Dotted line, Hb. White arrowheads, neurites from tdTomato^+^ HbX neurons extending into the MHb. **E,** Whole-mount immunostaining for tdTomato in adult *Cart-cre:Ai14* mice followed by 3DISCO and FLSM. Top: horizontal plane, bottom: coronal plane. Third ventricle, masked in yellow; Hb, masked in grey. Spots are color-coded per bin. Graphs: distance (x) of each spot from the third ventricle (i.e. along the medial-to-lateral axis). Dotted line, border MHb and LHb. N = 1 mouse/age. **F,** Immunohistochemistry for tdTomato, NRP2 (marks Hbcom) and/or PAX7 (marks IPN) in *Cart-cre:Ai14* mice. tdTomato^+^ fibers are observed in the Hbcom, FR and dorsal IPN. **G,** Immunohistochemistry for tdTomato and CHAT in a coronal section of adult *Cart-cre:Ai14* adult mice. Note tdTomato^+^/CHAT^+^ and tdTomato^+^/CHAT^-^ in the IPN. **H**, Location Dextran-488 tracer injection (yellow) in the dorso-lateral IPN. Purple, tdTomato immunohistochemistry. **I,** Representative image of Cartpt^+^/Dextran-488^+^ double-positive neurons (white arrows) in the ventral-lateral MHb following tracer injection as in H. Scale bar, 50 μm (I, lower panels), 200 μm. (A, B, F-I (upper)), 100 μm (C-E) See Figure S9.

To further establish the molecular identity of *Cartpt^+^* Hb neurons, the DevHb data were analyzed in more detail. This showed that *Cartpt* is expressed in the Hb from P4 onwards, as confirmed by ISH (Fig. 6B). *Cartpt^+^* neurons were found in multiple MHb clusters (most prominently in cluster 7 (SP-ergic (*Tac1^+^)*) and 9 (cholinergic (*Chat^+^* and also *Synpr^+^*)) that contributed to the border zone) and in the LHb cluster (11) (Fig. 6C, S5B). *Cartpt^+^* neurons of LHb cluster 11 (orange) contributed to the HbX and were marked by *Peg10* and *Gpr151* (Fig. 6C). Merger of the DevHb data with two adult Hb datasets, revealed *Cartpt* expression in adult MHb populations that correlate highly to DevHb clusters 7 and 9 (Fig. 2A, S9A-D).

Next, *Cartpt-cre:Ai14* mice (referred to here as *Cart-cre:Ai14*) were used to characterize projection patterns and functional properties of Hb *Cartpt^+^* neurons. Immunohistochemistry on sections of adult *Cart-cre:Ai14* mice showed that, in line with ISH data (Fig. 6A), tdTomato^+^ neurons locate to the HbX and border zone from P4 onwards (Fig. 6D). To establish their precise 3D localization, *Cart-cre:Ai14* mice were subjected to 3DISCO staining and FLSM. Quantification revealed that the majority of *Cartpt^+^* neurons is not only located at the border of the MHb and LHb, but also at the center of the Hb along the rostral-caudal axis (Fig. 6E, S9E). As the border zone contained the largest number of *Cartpt^+^* neurons, we focused on these cells. Most *Cartpt^+^* border zone neurons were *Chat^+^* and CHAT^+^ (Fig. 6C, S9F), similar to vMHb neurons. Since CHAT^+^ vMHb neurons project to the IPN (Albanese et al., 1985), we examined whether *Cartpt^+^* axons also target the IPN. Indeed, in P7 and adult *Cart-cre:Ai14* mice, tdTomato^+^ axons were detected in the IPN (Fig. 6F, S9G), and a subset of CHAT^+^ axons in the IPN was tdTomato^+^ (Fig. 6G). To unambiguously establish that *Cartpt^+^* neurons innervate the IPN, dextran-488 retrograde tracer was injected in the dorso-lateral IPN of adult *Cart-cre:Ai14* mice (Fig. 6H). At 7 days post-injection, a large number of tdTomato^+^/dextran-488^+^ neurons were detected in the border zone (Fig. 6H, I). Quantification showed that at least 35% of the total number of *Cartpt^+^* cells projects to the dorso-lateral IPN (n = 2 mice; N = 175 neurons). Conversely, dextran-488 injection into the VTA, a well-established projection site of LHb neurons, resulted in labeling of LHb, but not of the area of tdTomato^+^ neurons (Fig. S9H). Together, these results identify the dorso-lateral IPN as a major target of *Cartpt^+^* border zone neurons.

Although *Cartpt^+^* neurons share CHAT expression and projection targets with surrounding *Cartpt^-^* vMHb Hb neurons, they display a partially distinct molecular profile (e.g. *Cartpt*, *Sema3d* expression). To assess whether these molecular differences reflect distinct physiological properties, we performed whole-cell patch-clamp recordings using *Cart-cre:Ai14* mice (n = 13) (Fig. 7A). *Cartpt^+^* neurons (n = 17) had lower membrane capacitance than MHb (n = 15) and LHb (n = 10) *Cartpt^-^* cells (KW H(2) = 23.18, *P* = 0.000; *Cartpt^+^* vs. MHb *Cartpt^-^*, *P* = 0.017 and *Cartpt^+^* vs. LHb *Cartpt^-^*, *P* = 0.000)(Fig. 7B). This suggests that *Cartpt^+^* neurons are smaller than neighbouring MHb and LHb cells, which was confirmed by soma size measurements (1-way ANOVA F(2,33) = 8.95, *P* = 0.001; *Cartpt^+^* vs. MHb *Cartpt^-^, P* = 0.002 and *Cartpt^+^* vs. LHb *Cartpt^-^*, *P* = 0.001)(Fig. 7C). *Cartpt^+^* neurons also exhibited higher membrane resistance (1-way ANOVA F(2,41) = 6.46, *P* = 0.004; *Cartpt^+^* vs. MHb *Cartpt^-^, P* = 0.028 and *Cartpt^+^* vs. LHb *Cartpt^-^, P* = 0.001)(Fig. 7D). Potential differences in active conductance between *Cartpt^+^* and neighboring cells were also examined. *Cartpt^+^* neurons exhibited a higher spontaneous firing frequency, as indicated by the number of action potentials fired at rest (KW H(2) = 12.53, *P* = 0.002; *Cartpt^+^* vs. MHb *Cartpt^-^, P* = 0.149 and *Cartpt^+^* vs. LHb *Cartpt^-^, P* = 0.001). Instead, the action potential threshold was not significantly different (1-way ANOVA F(2,41) = 3.18, *P* = 0.052) (Fig. 7E, F). The DevHb dataset showed the presence of HCN channels in the *Cartpt^+^* cluster (Fig. S9I) and to assess whether the DevHb dataset could be used to predict the presence of active ionic conductance profiles, we measured the occurence of HCN-mediated voltage sags. *Cartpt^+^* cells indeed exhibited voltage sags, and these were more pronounced than in neighboring *Cartpt^-^* cells (KW H(2) = 8.65, *P* = 0.013; *Cartpt^+^* vs. MHb *Cartpt^-^, P* = 0.010 and *Cartpt^+^* vs. LHb *Cartpt^-^, P* = 0.372) (Fig. 7G). Together, these results establish that *Cartpt^+^* border zone neurons display a distinct electrophysiological profile, as compared to neighboring neurons. To assess how well one could predict whether or not a Hb neuron in the border zone belongs to the *Cartpt^+^* population, unbiased cluster analysis was performed based on cell capacitance and membrane resistance, the two variables that differed most between Cartpt^+^ cells and both MHb and LHb *Cartpt^-^* cells. This prediction yielded two independent clusters, in which 87.5% of Cartpt^+^ and 75% of *Cartpt^-^* cells fell into different clusters (Fig. 7H-J).

**Figure 7.**
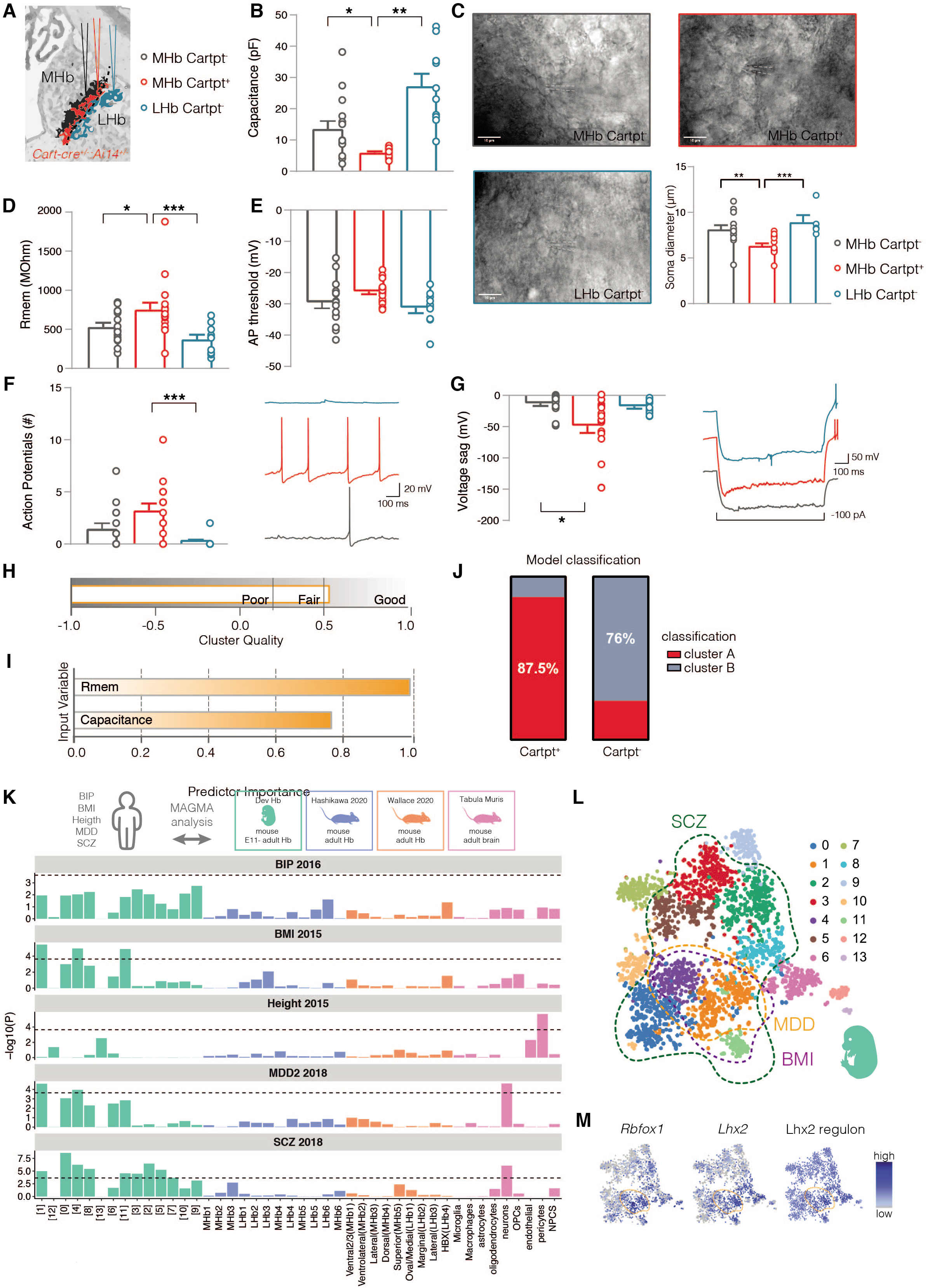
Cartpt^+^ neurons in the habenula border zone have distinctive functional properties and developing habenular subtypes associate with disease. **A,** Location of electrophysiological recordings in adult *Cart(pt)-cre:Ai14* mice (image adapted from Allen Brain Atlas). **B-G,** Cellular capacitance (pF) (B), cell soma Feret’s diameter (representative images are shown) (C), membrane resistance (Rmem) (D), AP threshold and number of spontaneous APs (E, F), and voltage sag (mV) (G) were determined in the three cell populations (see A). Representative traces are shown in F and G. Shown as mean ± S.E.M. * P<0.05, ** P<0.01, *** P<0.001. **H-J,** Recorded cells were classified by a two-step cluster analysis of individual data on cell capacitance and Rmem using the Schwarz’s Bayesian criterion. Silhouette measure of cohesion and separation revealed an overall good model fit (average silhouette 0.5). Rmem was the most important predictor of cell clustering (I). 87.5% of all Cartpt^+^ and 76% of all Carpt^-^ neurons were classified in accordance to their genotype based on capacitance and Rmem. **K,** MAGMA analysis, cell types are color-coded by dataset. Bar plots indicate signal strength, if dashed line (*P* value after Bonferroni correction = 0.05 / (44*5)) is crossed, signal is considered significant. **L,** t-SNE embedding color-coded by Louvain cluster. Associated clusters per trait are highlighted by dashed lines. **M,** t-SNE embedding showing the relative expression levels of MDD risk genes *Rbfox1* and *Lhx2*, and *Lhx2* regulon activity. See Figure S9, 10.

Overall, these data provide insight into the development and molecular identity of *Cartpt^+^* Hb neurons, unveil a major projection target of these neurons, and show that Cartpt^+^ neurons represent a functionally distinct population in the Hb border zone.

### Developing habenular subtypes align to GWAS-identified risk loci of human disease

Finally, we assessed whether our scRNAseq data could aid in the identification of cellular subtypes related to Hb-associated diseases, such as psychiatric disorders. By performing MAGMA gene-property analysis, in which gene expression levels per cluster are gene-properties (de Leeuw et al., 2015; Watanabe et al., 2019), the convergence of risk variants of a specific trait onto a specific cell type was investigated. For this analysis psychiatric traits were selected for which involvement of the Hb had been implied (MDD, schizophrenia (SCZ) and bipolar disorder (BIP) (Hou et al., 2016; Ripke et al., 2014; Wray et al., 2018). Given the involvement of the Hb in feeding behavior, body mass index (BMI) was also included (Locke et al., 2015). Height was included as a control, as no association was suspected (Wood et al., 2014) (Fig. 7K). To confirm the validity of our approach, the Tabula Muris dataset was used to show that neurons are implicated in MDD and SCZ, as shown previously (Watanabe et al., 2019) (Fig. 7K). Next developmental Hb cell types (DevHb data) and adult Hb cell types, were analyzed. BIP showed no association with the selected clusters, and height only associated with pericytes from the Tabula Muris dataset. BMI showed an association with DevHb clusters 1, 4 (iHb) and 11 (LHb)(Fig. 7K, L). Interestingly, clusters composed only of adult Hb neuron subtypes did not associate with any of the tested traits and diseases. In contrast, MDD and SCZ were associated with DevHb clusters 1 and 4, and SCZ in addition with clusters 0, 2, 3, 5, 8, 11 (Fig. 7K, L). The known MDD risk genes *Rbfox1* and *Lhx2* (Wray et al., 2018) were highly expressed in clusters 1 and 4, which contain iHb that will mature into *Calb2*^+^ dMHb neurons and LHb neurons (Fig. 7M). Further, expression of the *Lhx2* regulon was particularly high at the bridge between cluster 6 (no MDD association) and cluster 1 (MDD association) (Fig. 7M). Next, we asked whether the detected associations with MDD and SCZ are specific to developing Hb neuron subtypes. Two additional datasets, containing developing hypothalamus neurons (Kim et al., 2020) or developing midbrain dopamine neurons (La Manno et al., 2016), were randomly down-sampled and analyzed. SCZ associated with developing midbrain dopamine and hypothalamus neuron subsets, and for MDD an association with developing hypothalamic neurons was found (Fig. S10).

Overall, this analysis shows that both MDD and SCZ are associated with specific developmental Hb subtypes, and to a varying degree with other developing neuronal subtypes.

## DISCUSSION

Hb neurons have complex connectivity patterns and functions, and their disruption is linked to various disorders. The mouse Hb is composed of several different subtypes of neurons, but how these subtypes develop, connect and function remains poorly understood. Progress in this regard is hampered by a lack of knowledge of the developmental gene expression profiles and trajectories of these neuronal subtypes. To address this void, we have combined scRNAseq with molecular, anatomical and functional approaches. Our data reveal novel cellular and molecular trajectories during development leading to different adult Hb subtypes. Further, our work establishes the projection target and distinctive functional properties of a previously uncharacterized subtype of *Cartpt^+^* Hb border zone neurons. Finally, comparison of developing Hb clusters and GWAS data suggests that subtypes of developing Hb neurons associate with psychiatric disorders (Fig. S10G). Our study begins to dissect the molecular and cellular basis of Hb neuron subtype development and constitutes a valuable resource for future studies on Hb development in health and disease.

### Cellular diversity in the developing mouse habenula

Previous studies have identified molecularly distinct neuron subtypes in the adult Hb (e.g. (Aizawa et al., 2012; Amo et al., 2010; Andres et al., 1999; Hashikawa et al., 2020; Le Foll and French, 2018; Pandey et al., 2018; Wagner et al., 2016; Wallace et al., 2020)). However, the ontogeny, projection targets, functional properties and disease-associated roles of most of these subtypes are unknown. To address this key unresolved question, we performed scRNAseq on Hb cells sorted from *Brn3a-tauLacZ* mice at multiple developmental stages. *Pou4f1/Brn3a* is an accepted marker for post-mitotic Hb neurons (Guo and Li, 2019; Quina et al., 2010) and labels ±80 % of MHb and ±55 % of LHb neurons (Fig. S2B). In line with these numbers, trajectory endpoints of MHb populations in the DevHb data matched previously defined adult MHb populations, whereas cellular heterogeneity in the LHb was not fully captured (one LHb cluster (11) correlated to the multiple reported adult LHb clusters). Nevertheless, *Brn3a-tauLacZ* mice are currently the most efficient tool for purifying Hb neurons throughout development.

Analysis of the DevHb and E18 WT data identified a large cluster of *Id3^+^/Vim^+^* PCs. BrdU birth dating and neurosphere assays confirmed a population of PCs in the E18 Hb ventricular zone (Fig. S10E, F). *Id3^+^/Vim^+^* PCs developed into iHb neurons expressing high levels of *Cntn2, Dcc* and *Flrt3*. Then the trajectory split into separate axes of the LHb, dMHb and vMHb. This suggests that mouse LHb and MHb neurons originate from the same PCs. This is interesting as equivalent zebrafish structures derive from distinct PC populations (Beretta et al., 2013). A high level of cellular heterogeneity was already detected as early as E11 in our data, but mouse Hb subtypes with adult signatures were only present by P4. Therefore, our data for the first time establish that Hb subtypes acquire adult transcriptional signatures only at early postnatal stages.

### Molecular mechanisms underlying habenula development

Different aspects of Hb development have been studied (e.g. (Aizawa et al., 2007; Beretta et al., 2013; Chatterjee et al., 2014; Colombo et al., 2013; Fore et al., 2020; Guglielmi et al., 2020; Hüsken et al., 2014; Kantor et al., 2004; Quina et al., 2009; Ruiz-reig et al., 2019; Schmidt et al., 2014)), but how specific neuronal subtypes develop remains largely unexplored, especially in mammalian species. As highlighted below, our study begins to address the mechanisms underlying Hb subtype-specific development. For example, we report developmental stage-specific expression or enrichment of axon guidance proteins and receptors (and other cues) in select developmental clusters. This was especially prominent for EPHRINS and EPH receptors, a large family of axon guidance proteins involved in topographic mapping of axons (Kania and Klein, 2016). This is interesting as neurons in different Hb subdomains target specific brain (sub)nuclei (Herkenham and Nauta, 1977; Quina et al., 2015; Wallace et al., 2020) and because subdomain-specific EPHRIN/EPH expression is found in these nuclei (García-Guillén et al., 2021). These molecular insights will aid the future dissection of mechanisms instructing Hb subtype-specific circuit formation.

Previous studies unveiled Hb subtype-specific expression of genes associated with synaptic transmission at adult stages (e.g. (Pandey et al., 2018; Wallace et al., 2020)). Our data allowed spatiotemporal analysis of the expression of these genes. Expression of ‘synaptic transmission’ genes was generally first detected at postnatal stages, although striking differences were observed. For example, *Tac1* (marker of SP-ergic neurons) was found in the early embryonic dMHb, while *Chat* (marker of cholinergic neurons) was first detected at postnatal stages in the vMHb. Future studies are needed to establish whether *Tac1^+^* neurons are born earlier than *Chat^+^* neurons, or whether these populations only differ in their onset of gene expression.

Our work also identified different developmental trajectories leading to mature Hb neuron subtypes. These were marked by trajectory-specific gene expression trends which may contribute to the transition of PCs into MHb and LHb neuron subtypes. To better understand the molecular basis of neuron subtype-specific differentiation, we attempted to identify specific TF-driven gene regulatory networks. Differences in these networks were observed between developmental stages but not Hb subtypes (except for PCs). Nevertheless, our results provide novel and detailed insight into the developmental stage-specific gene regulatory networks underlying Hb development.

### Linking molecular and functional properties of habenular subtypes

Whether molecularly defined Hb subtypes have distinctive functional properties or connections is incompletely understood. Here, we characterized *Cartpt^+^* neurons that mark the HbX and the border zone, an area at the boundary of MHb and LHb (Wagner et al., 2016; Wallace et al., 2020). *Cartpt* encodes for cocaine- and amphetamine-regulated transcript (CART) peptide which affects locomotor activity and has anti-depressant effects (Singh et al., 2021). However, *Cartpt*^+^ Hb neurons, and HbX and border zone neurons in general, remain poorly characterized. Using immunohistochemistry and genetic tools *Cartpt*^+^ Hb neurons were detected from early postnatal stages onwards. Interestingly, transcripts displaying an expression pattern similar to *Cartpt* included *Sema3d* (Wagner et al., 2016; Wallace et al., 2020). SEMA3D is a member of the semaphorin family of guidance proteins with roles in axon guidance and synaptogenesis (Pasterkamp, 2012) and may play a role in regulating *Cartpt^+^* Hb neuron connectivity. For example, in establishing highly distinctive patterns of afferent innervation at the border zone (Herkenham and Nauta, 1977; Shinoda and Tohyama, 1987).

Analysis of *Cart-cre:Ai14* mice and retrograde tracing showed that many *Cartpt^+^* border zone neurons express CHAT and send their axons to the IPN, both characteristics of MHb neurons. However, despite these similarities, distinctive electrophysiological profiles were found for *Cartpt^+^* neurons as compared to surrounding *Cartpt^-^* MHb and LHb neurons. Differences in cell capacitance and membrane resistance hinted at distinct morphology and indeed cell body diameter measurements confirmed a significantly smaller soma size for *Cartpt^+^* as compared to surrounding *Cartpt^-^* neurons. Higher membrane resistance indicates that *Cartpt^+^* neurons are more electrotonically compact, which could be mediated by differential potassium channel expression (Tripathy et al., 2017). Finally, *Cartpt^+^* neurons displayed prominent voltage sag, in accordance with predictions from the DevHb dataset regarding the presence *Hcn3* and *Hcn4* ion channels in *Cartpt^+^* cell clusters. Together, our data show that *Cartpt^+^* border zone neurons represent a molecularly and functionally distinct subtype of Hb neurons that send prominent axon projections to the IPN. The functional role of this circuitry is unknown, but the IPN mediates reward processing and depressive-like symptoms (Xu et al., 2018), in line with reported roles of CART peptide (Ahmadian-Moghadam et al., 2018).

### A role for developing habenular subtypes in human disease?

Hb dysfunction is linked to depression (Hu et al., 2018) and GWAS of human MDD patients uncovered multiple genetic risk loci (Wray et al., 2018). MAGMA analysis of our data, and other published datasets, revealed a significant association between MDD risk variants and developing Hb neurons. In the associated subtypes high expression of the MDD risk genes *Rbfox1* and *Lhx2* was observed. Interestingly, *Rbfox1* was especially high in young Hb neurons that eventually mature into LHb and *Calb2^+^* dMHb neurons. These results unveil a previously unreported link between the developing Hb and MDD and hint at the possibility that changes in the development of LHb and *Calb2^+^* dMHb neurons contribute to the etiology of this psychiatric disorder. Immature Hb populations, and other types of developing neurons, also associated with SCZ risk genes. This is in line with previous work connecting SCZ and embryonic neuroblasts (Watanabe et al., 2019), but contrasts data showing lack of association between SCZ and developing neurons (Skene et al., 2018). Thus, although further work is needed to address these apparent discrepancies, our proof-of-concept analysis generates an interesting framework for future studies on the link of the developing Hb and disease.

In summary, we have harnessed the power of scRNAseq, in combination with other techniques, to dissect the developmental programs underlying Hb neuron diversity. Our analysis provides examples of cell state markers, developmental trajectories, and transcriptional programs. Further, it also contributes a unique 3D overview of Hb development and the functional characterization of an undefined molecular subtype of Hb neurons. Finally, it reveals that comparison of Hb scRNAseq and disease GWAS data may provide insight into disease pathogenesis (Fig. S10G). Together, our data provide an initial framework for dissecting the developmental programs underlying Hb neuron diversity and a starting point for future interrogation of the developing Hb in normal and disease states.

## ACKNOWLEDGEMENTS

We thank Dinos Meletis for input on our manuscript and members of the Pasterkamp lab for help and discussions; Eric Turner for *Brn3a-tauLacZ* mice; Nefeli Kakava for technical help; Kiochi Hashikawa, Garret Stuber, Michael Wallace, Bernardo Sabatini, Shristi Pandey and Alexander Schier for sharing data. This research was supported by the Netherlands Organization for Scientific Research (ALW-VICI 865.14.004 to R.J.P.), ENW-VENI (016.Veni.192.188 to D.R.), ERC under the European Union’s Horizon 2020 research and innovation programme (804089; ReCoDe to F.J.M.). Partially supported by NWO Gravitation program BRAINSCAPES: Roadmap from Neurogenetics to Neurobiology (NWO: 024.004.012), Stichting Parkinson Fonds (to R.J.P.).

## AUTHOR CONTRIBUTIONS

L.L.vd.H. and R.J.P. designed the study and wrote the manuscript with help from all authors. L.L.vd.H. and D.R. designed and performed experiments with help from J.E.B., T.E.S., R.E.v.D., Y.A., C.V.D.M., N.C.H.v.K.; and M.H.B. D.P. and K.W. helped with performing MAGMA analysis. F.J.M. and O.B. designed experiments and aided in data analysis.

## STAR METHODS

### RESOURCE AVAILABILITY

#### Lead Contact

Further information and requests for resources and reagents should be directed to and will be fulfilled by the Lead Contact, Jeroen Pasterkamp.

#### Materials Availability

All unique reagents, plasmids, and transgenic mouse lines generated in this study are available from the Lead Contact with a completed Materials Transfer Agreement.

#### Data and Code Availability

All data is available through GEO: GSE188712. All code used to analyze the scRNAseq data can be found at https://github.com/liekevandehaar/Dev_Hb_mouse

### EXPERIMENTAL MODEL AND SUBJECT DETAILS

#### Mouse lines

All animal experiments in this study were approved by the (CCD) Centrale Commissie Dierproeven of Utrecht University (CCD license: AVD115002016532) and were in accordance with Dutch law (Wet op de Dierproeven 2014) and European regulations (guideline 2010/63/EU). *Brn3a-tauLacZ* (Raisa Eng et al., 2001) (gift from prof. Eric Turner), *Cart-IRES2-cre-D* (The Jackson Laboratory, JAX:028533), *STOP-tdTomato (Ai14)* (The Jackson Laboratory, JAX:007914) and *C57BL/6j* mice (Charles Rivers) were housed at 22 ± 1°C on a wood-chip bedding supplemented with tissue on a 12 hr/12 hr day/night cycle. Pregnant mothers were housed individually from the moment of observation of the plug (E0.5). Animals were fed *ad libitum* and mice from both sexes were used. Frozen sperm of *Brn3a-tauLacZ* males was provided by prof. Eric Turner. Animals were generated as described (Raisa Eng et al., 2001). In short, a tauLacZ reporter cassette was inserted in part of the first and second exon of the *Brn3a* locus. Frozen sperm was recovered and viable mice were generated at Janvier Labs (France). Mice were selected for the presence of the *Brn3a-tauLacZ* locus. Because homozygous animals die shortly after birth, animals were kept heterozygous on a *C57BL/6j* background. Animals were genotyped using the primers listed in Table S4.

#### Neurosphere generation

Cells were isolated as described previously (Guo et al., 2012a). Adult mice were sacrificed by cervical dislocation. Brains were isolated and placed in ice-cold 1x PBS. The Hb and the subventricular zone (SVZ) were dissected in ice-cold L15++ medium (L15 (Thermo-Fisher, Cat#11415049) supplemented with 7.5 mM HEPES (Thermo-Fisher Cat#15630106) and 1 x penicillin/streptomycin (pen/strep; Thermo-Fisher, Cat# 15140122)). Tissue was dissociated using the MACS Neural Dissociation kit (Miltenyi Biotec, Cat#130-093-231) according to the manufacturer’s protocol. In short, tissue was incubated in pre-heated dissociation solution (200 μl Trypsin and 1750 μl solution X) at 37°C for 15 min. 20 μl solution Y and 10 μl solution A were added per sample. Tissue was dissociated with a wide-tipped glass pipet. The samples were incubated at 37 °C for 10 min. Tissue was further dissociated with a series of 2 glass pipettes with descending tip sizes. 5 ml glutaMAX DMEM/F-12 (Thermo-Fisher, Cat#10565018) supplemented with 1x pen/strep was added to each sample. The suspension was filtered using a 40 μm cell strainer into a 15 ml Falcon tube and centrifuged at 300 x g for 5 min. The cell pellet was washed with L15++ medium and centrifuged at 300 x g for 5 min. Cells were resuspended in glutaMAX DMEM/F-12 (Thermo-Fisher, Cat#10565018) supplemented with 20□g/ml EGF (Thermo-Fisher, Cat#PHG0311L), 20□ng/ml FGF (Thermo-Fisher, Cat#PGH0367), B-27 (Thermo-Fisher, #17504044) and pen/strep (Thermo-Fisher, Cat#15140148) and kept in a 5% CO_2_ incubator at 37°C.

### METHOD DETAILS

#### Sample collection for scRNAseq

Pregnant *Brn3a-tLacZ^+/-^* or *C57BL/6j* mice were sacrificed by cervical dislocation. Embryos were harvested at embryonic (E) day E11, E12, E13, E15, and E18. The uterine horn was placed in ice-cold 1x PBS. Embryonic brains were dissected in ice-cold medium A (1x HBSS – Ca^2+^ - Mg^2+^ (Thermo-Fisher, Cat#24020117)) supplemented with 1.6 mM HEPES (Thermo-Fisher, Cat#15630106) and 0.63% glucose. Tissue was dissociated using the MACS Neural Dissociation kit (Miltenyi Biotec, Cat#130-093-231) for 30 min according to the manufacturer’s protocol. Postnatal (P4 – P7) and adult *Brn3a-tLacZ^+/-^* mice were cervically dislocated (only adult: P245-P259) and decapitated. The heads were stored on ice. The brain was removed from the skull and placed in ice-cold HABG medium (Hibernate A – Ca^2+^ - Mg^2+^ (BrainBits, Cat#HACAMG500)) supplemented with 1x B27 (Thermo-Fisher, Cat#17504044) and 0.5 mM L-glutamine (Thermo-Fisher, Cat#25030081)). The Hb was dissected out, manually cut into smaller pieces and placed in 2 ml HABG medium in a 15 ml Falcon tube on ice. Samples were spun down at 1000 rpm for 1 min. Supernatant was taken and 0.5 ml warm papain diluted in HABG was added. Samples were broken by pipetting 5 times with a P1000 pipet. Samples were incubated horizontally on a shaker for 15 min at 160 RPM at 37°C. 35 μl DNase mix 1 (1 volume enzyme X (DNase) (Neural Dissociation kit (T) (Miltenyi Biotec, Cat#130-093-231)) and 2 volumes buffer Y (Neural Dissociation kit (T) (Miltenyi Biotec, Cat#130-093-231)) was added to the sample. Next, the sample was incubated horizontally on a shaker for 15 min at 160 RPM at 37°C. 135 μl warm DNase mix 2 (30 μl DNase mix 1, 100 μl FBS, 2 μl BSA 1 mg/ml (Jackson, Cat#001-00-162) and 1 μl 0.5 M EDTA) was added to each sample. For complete dissociation, samples were triturated 2 – 3 times with 3 different glass pipettes decreasing in size. Samples were passed through a 100 μm filter and 5 ml aCSF (92 mM NaCl, 2.5 mM KCl, 1.2 mM NaH_2_PO_4_, 30 mM NaHCO3, 20 mM HEPES (Thermo-Fisher, Cat#15630106)), 25 mM glucose, 3 mM sodium ascorbate, 2 mM thiourea and 3 mM sodium pyruvate) supplemented with 10% FBS, and 5 mM EDTA was added. Samples were centrifuged at 100 g for 10 min. Supernatant was removed, cells were taken up in 100 μl staining medium (4% FBS and 10 mM HEPES (Thermo-Fisher, Cat# 15630106), in 1x PBS), filtered through a 70 μm FACS filter, and placed on ice.

FluorLacZ staining was performed using the FluoReporter lacZ Flow Cytometry Kit (Thermo Fisher Scientific, F1930) according to the manufacturer’s protocol. Cell in staining medium and the FDG (fluorescein di-β-D-galactopyranoside) working solution (2 nM FDG in MQ) were pre-warmed for 10 min at 37°C. 100 μl of FDG working solution was added to the cells and rapidly mixed. Cells were placed in a water bath at 37°C for exactly 60 sec. To stop the FDG loading 1.8 ml ice-cold staining medium containing 1.5 μM propidium iodide was added to the cells using ice-cold pipettes and cells were stored on ice.

Cells were sorted into 384-wells plates in a BD FACS ARIAII FACS machine with a 100 μm nozzle. Cell were sorted based on DAPI-negativity (E18 WT experiment) and DAPI negativity followed by LacZ positivity (*Brn3a-tLacZ* developmental experiment) (Figure S2C, D). Each replicate consists of two 384-well plates each containing an individual sample. After sorting, cells were spun down at 1000 rpm for 1 min, placed on dry ice and stored at −80°C.

#### scRNAseq sequencing

Cells were sequenced by Single Cell Discoveries (Utrecht, The Netherlands) according to the SORT-seq method based on CEL-Seq2 (Muraro et al., 2016). In short, cells were lysed and the poly-T stretch of the barcoded primers hybridized to the poly-A tail of the mRNA molecules. The DNA-RNA hybrids were converted into cDNA containing the mRNA and primer sequences. All cDNA was pooled and amplified by *in vitro* transcription. Finally, PCR selected for molecules that contain both Illumina adaptors necessary for sequencing. Subsequently, bead cleanup and quality control were performed with the Agilent High Sensitive DNA Kit (5067-4626) and libraries were sequenced by the Utrecht Sequencing Facility (USEQ, Life Sciences Faculty, Utrecht University). Paired end sequencing was performed by Nexseq500.

#### 3DISCO

3DISCO was performed as described previously (Adolfs et al., 2021; Belle et al., 2014). In short, E10, E11, E12 and E13 whole mouse embryos, and E15 and E18 mouse brains were placed in 4% PFA, pH 7.4, for 24 hr at 4°C. Samples were washed in 1x PBS for ≥ 1.5 hr on a shaker at room temperature (RT). Subsequently, they were dehydrated using a series of 50%, 80% and 100% MeOH (Merck Millipore, Cat#1060092500), each step was performed for ≥ 1.5 hr on a shaker at RT. Samples were bleached overnight at 4°C without shaking in 3% hydrogen peroxide (Merck Millipore, Cat# 1072091000) in MeOH. The following day samples were hydrated using a series of 100%, 100%, 80%, 50% MeOH and 1x PBS for ≥ 1 hr on a shaker at RT. The bleached samples were placed in a 15 ml Falcon tube containing blocking buffer (PBSGT) consisting of 1x PBS with 0.2% gelatin, 0.5% Triton-X-100 (Sigma, Cat#x 100-500ml), and 0.01% thimerosal (Gerbu Cat#1031/USP35) at RT on a horizontal shaker at 70 RPM. Incubation time varied from 3 hr (E10-E12) to 24 hr (E13-E18). Samples were incubated at 37°C on a horizontal shaker (70 RPM) in a 2 ml Eppendorf tube containing 2 ml PBSGT, 0.1% saponin (Sigma, Cat#S7900), and primary antibody.

Antibodies used were goat anti-Neuropilin-2 (NRP2) (R&D Systems, Cat#AF567, 1:1000), rabbit anti-Tyrosine Hydroxylase (EMD Millipore, Cat#TH152, 1:300), goat anti-ROBO1 (R&D Systems, Cat#AF1749, 1:300), and goat anti-ROBO3 (R&D Systems, Cat#AF3076, 1:300). Incubation times varied from 3 days (E10-E13) to 7 days (E15-E18). Samples were washed 6 times for 1 hr at RT in a 15ml Falcon tube containing PBSGT on a rotator at 14 RPM. Next, samples were incubated at 37°C for 48 hr on a horizontal shaker (70 RPM) in a 2 ml Eppendorf tube containing 2 ml PBSGT, 0.1% saponin (Sigma, Cat#S7900) and secondary antibody protected from the light from this point onwards. Antibodies used were Alexa Fluor 647 donkey anti-goat (Abcam, Cat#ab150135, 1:500) and Alexa Fluor 750 donkey anti-rabbit (Abcam, Cat#ab175731, 1:500). Next, samples were washed 6 times for 1 hr at RT in a 15 ml Falcon tube containing PBSGT on a rotator at 14 RPM. After staining, E10 and E11 embryos were embedded in 1% agarose dissolved in 1x PBS. For clearing, tissue samples were placed in a series of 50% (overnight), 80% (1 hr), 100% (1 hr) and 100% (1 hr) tetrahydroflurane with BHT as inhibitor (Sigma, Cat#186562-1L). All steps were performed in a 15 ml Falcon at RT on a horizontal shaker at 70 RPM. To dissolve lipids, samples were placed in 100% dichloromethane (DCM) (Sigma, Cat#270997-1L) for 20 min or until they sank. Finally, samples were incubated overnight in 100% dibenzyl ether (DBE) (Sigma, Cat#108014-1kg) in a 15 ml Falcon tube at RT on a rotator at 14 RPM. Samples were stored in 100% DBE at RT protected from light. Samples were imaged using an Ultramicroscope II (LaVision BioTec) light sheet microscope equipped with an MVX-10 Zoom Body (Olympus), MVPLAPO 2x Objective lens (Olympus), Neo sCMOS camera (Andor) (2560 x 2160 pixels. Pixel size: 6.5 x 6.5 μm^2^) and Imspector software (version 5.0285.0, LaVision BioTec). Samples were scanned with single-sided illumination, using the horizontal focusing light sheet scanning method and the blend algorithm. The Object lens included a dipping cap correction lens (LV OM DCC20) with a working distance of 5.7 mm. Imaris (version 8.4-9.4, Bitplane) software was used for image processing and analysis.

#### iDISCO

*Cart-IRES2-cre-D^+/^:Ai14^+/-^* mice (E15, P4, P7 and adult) were stained using the iDISCO method described previously (Renier et al., 2016). In brief, samples were dehydrated in a 15 ml Falcon tube containing a series of 20% MeOH, 40% MeOH, 60% MeOH, 80% MeOH, or 100% MeOH, each for 1 hr at RT followed by overnight incubation at RT in 66% DCM and 33% MeOH. The following day samples were bleached by two times incubation at RT in 100% MeOH, followed by incubation of the samples at 4°C for 3 hr. This was followed by incubation in 2% hydrogen peroxide in MeOH overnight at 4°C. The following day samples were rehydrated using a series of 80% MeOH, 60% MeOH, 40% MeOH, 20% MeOH, each for 1 hr at RT. This was followed by washes in 1x PBS, 1 hr at 4°C. Next, samples were incubated twice in Ptx.2 (1x PBS and 0.2% Triton-X100) for 1 hr at RT and placed in permeabilization solution for 24 hr at 37°C. Then, samples were incubated in blocking solution for 4 days at 37°C on a horizontal shaker (70 RPM). Subsequently, samples were incubated in primary antibody solution (PwtH (1 x PBS, 0.2% Triton-X100, 0.1% Heparin (Sigma, Cat#H3393, 10 mg/ml) stock solution, 5% DMSO and 3% normal donkey serum (Jackson Immunoresearch, Cat#017-000-121)) for 7 days at 37°C on a horizontal shaker (70 RPM). Then, samples were washed 5 times for 1 hr at RT in a 15 ml Falcon tube containing PtwH. Samples were incubated in secondary antibody solution (PwtH and 3% normal donkey serum (Jackson Immunoresearch, Cat#017-000-121)) for 7 days at 37°C on a horizontal shaker (70 RPM) in a 5 ml Eppendorf. Then, samples were dehydrated using a series of 20% MeOH, 40% MeOH, 60% MeOH, 80% MeOH and 2 times 100% MeOH, each step for 1 hr at RT. Then, samples were placed in 66%DCM and 33% MeOH for 3 hr at RT, followed by 2 washes in 100% DCM for 15 min at RT each. For clearing, tissue samples were placed in 100% DBE overnight at RT. Samples were stored in 100% DBE at RT until imaging. All washing, dehydration and clearing steps were performed in dark Falcon tubes to protect against light and on a rotator (14 RPM). Primary antibodies used were rabbit anti-RFP (Rockland, Cat#600-401-379, 1:1000) and goat anti-ROBO3 (R&D, AF3076, 1:300). Secondary antibodies used are Alexa Fluor 647 donkey anti-rabbit (Abcam, Cat#ab150135, 1:1000) and Alexa Fluor 568 donkey anti-goat (Abcam, Cat#ab175704, 1:1000). Samples were imaged using an Ultramicroscope II (LaVision BioTec) light sheet microscope equipped with an MVX-10 Zoom Body (Olympus), MVPLAPO 2x Objective lens (Olympus), Neo sCMOS camera (Andor) (2560 x 2160 pixels. Pixel size: 6.5 x 6.5 μm^2^) and Imspector software (version 5.0285.0, LaVision BioTec). Samples were scanned using single-sided illumination, a sheet NA of 0.148 and a step-size of 2.5 μm using the horizontal focusing light sheet scanning method with 200 steps and using the blend algorithm. The Object lens included a dipping cap correction lens (LV OM DCC20) with a working distance of 5.7 mm. The effective magnification was 8.4208 x (zoom body*objective + dipping lens = 4 * 2.1052). Imaris (version 8.4-9.4, Bitplane) software was used for image processing and analysis.

#### Whole-mount X-gal staining

The X-gal staining protocol was based on (Wythe et al., 2013). Embryos were fixed for 24 hr in fixative solution containing 2% formaldehyde (Riedel-de Hain, Cat#33200), 0.2% glutaraldehyde (Acros Organics, Cat# 119989925), 0.02% sodium deoxycholate (Sigma, Cat#D6750-100G) and 0.01% NP-40 in 1 xPBS. Samples were rinsed 2 times with ice-cold 1x PBS to remove fixative. Samples were permeabilized overnight at 4°C while shaking gently in permeabilization solution containing 0.02% sodium deoxycholate and 0.01% NP-40 in 1x PBS. After permeabilization samples were incubated in staining solution containing 1 mg/ml X-gal, 5 mM K-ferricyanide (Merck, Cat#231-847-6), 5 mM K-ferrocyanide (Sigma, Cat#p8131-100G), 2 mM MgCL_2_ (Merck, Cat#A0344733339), 0.02% sodium deoxycholate and 0.01% NP-40 in 1x PBS. Incubation was performed at 37°C and in the dark for 3 to 24 hr while shaking. Samples were washed 2 times for 15 min with permeabilization solution to remove staining solution. Embryos were dehydrated in a series of 30%, 50%, 80%, 100% and 100% EtOH. For clearing, embryos were incubated overnight at RT in 100% dibenzyl ether (DBE) (Sigma, Cat#108014-1kg) in a 15 ml Falcon tube on a horizontal shaker at 70 RPM. Embryos were imaged using a Zeiss Primovert inverted microscope using Zen 2 blue edition software (version 2.0.0.0).

#### In situ hybridization

##### Probe generation

Primers (Table S4) were designed to amplify the desired probe sequence from total mouse cDNA. PCR products were run in a 1.5% agarose gel and the correct amplicon was purified using the PureLink Quick Gel Extraction kit (Thermo-Fisher Cat#K210025). The purified DNA fragment was ligated into the pGEM-T-easy vector (Promega, Cat#A1360) according to the manufacturer’s protocol. Ligation products were transformed into DH5-α cells (Invitrogen, Cat#: 18263012). Upon confirmation by restriction digestion, inserts were sequenced using T7 and SP6 primers. The PCR product was used for SP6/T7 reverse transcription to generate DIG-labeled sense and anti-sense probes as described previously (Brignani et al., 2020). Probes were treated with 2 units of DNase (Roche, Cat#776785) for 15 min at 37°C. Then probes were centrifuged briefly and a solution containing 2 μl of 2.0 M EDTA, pH 8.0 (Sigma, Cat#E5134-500G), 2.5 μl of 4 M LiCl (Merck, Cat#7447-41-8), and 75 μl pre-chilled 100% EtOH was added. Samples were placed for 30 min at −80°C followed by centrifugation at 14,000 RPM at 4°C for 15 min. The pellet was washed with 50 μl prechilled 70% EtOH and centrifuged at 14,000 RPM at 4°C for 5 min. The pellet was air-dried for 10 min at RT, resuspended in 100 μl Milli-Q water, and stored at −80°C.

##### In situ hybridization

*In situ* hybridization was performed as described previously (Brignani et al., 2020). In brief, sections were defrosted and dried for 1.5 hr at RT. Subsequently, sections were fixed for 10 min in 4% PFA (Sigma #P-6148) in 1x PBS, pH 7.4. After fixation, sections were washed 3 times for 5 min in 1x PBS. Sections were placed in acetylation solution (1.32 % triethanolamine (Fluka Cat#90279), 0.18% HCl and 0.25% acetic anhydride (Sigma Cat#A6404)) for 10 min. After acetylation, sections were washed 3 times for 5 min in 1x PBS. Sections were pre-hybridized for 2 hr at RT in hybridization solution (50% deionized formamide (ICN, Cat#800686), 5x saline-sodium citrate (SSC; 0.75 M NaCl, 75 mM NaCitrate, pH 7.0), 5x Denhardts (1 mg/ml Ficoll-400 (Sigma, Cat#F4375)), 1 mg/ml polyvinylpyrrolidone (Merck, Cat#7443), 1 mg/ml BSA-fraction V (ICN Cat#103703), 250 μg/ml tRNA baker’s yeast (Sigma, Cat#R-6750), and 50 μg/ml sonificated Salmon Sperm DNA (Sigma, Cat#D-91565) in MQ). DIG-RNA probe was diluted in 150 μl hybridization solution and added to the sections. Sections were covered with NESCO-film and hybridized in a humidified chamber overnight at 68°C. To remove non-bound probe, slides were dipped in 2x SCC (0.3 M NaCl, 30 mM NaCitrate, pH 7.0) at 68°C and then washed in 0.2x SCC (30 mM NaCl, 3 mM NaCitrate, pH 7.0) for 2 hr at 68°C. Slides were transferred to RT 0.2x SSC and washed for 5 min. Slides were then washed for 5 min in buffer 1 (100 mM TrisHCl, pH 7.4, 150 mM NaCl) at RT. Sections were incubated for 1 hr at RT in buffer 1 supplemented with 10% FCS followed by incubation with anti-DIG-AP primary antibody (Sigma, Cat#11207733910, 1:5000) in buffer 1 containing 1% FCS overnight at 4°C. The following day, sections were washed 3 times for 5 min in buffer 1 and once for 5 min in buffer 2 (100 mM TrisHCl, pH 9.5, 50 mM MgCl2, 100 mM NaCl). Slides were developed in filtered developing solution (NBT/BCIP 1:20 (Boehringer Mannheim, Cat#1681451) and 0.24 mg/ml Levamisol (Sigma, Cat#L9756) in buffer 2). Slides were developed for 2 to 24 hr. Sections were mounted with glycerol. Images were obtained using an Axio Imager M2 microscope (Zeiss) and processed in Fiji (Schindelin et al., 2012) (version 2.0.0-rc-69/1.52o).

#### Stereotactic injections

For retrograde anatomical studies, male and female *Cart-IRES2-cre-D^+/^:Ai14^+/-^* mice (12-14 weeks old) were injected with 0.2-0.5 μl Alexa Fluor 488-conjugated Dextran (Invitrogen, diluted 1:3 in sterile PBS). Prior to surgery, mice were anaesthetized by intraperitoneal (i.p.) injection of ketamine (Narketan 10, Vétoquinol, 75 mg/kg) and medetomidine (Sedastart, Ast Farma B.V., 1 mg/kg). Lidocaine (B. Braun, 0.1 ml, 10% in saline) was administered subcutaneously under the scalp to provide local anaesthesia. Eye ointment (CAF, Ceva Sante Animale B.V.) was applied before surgery. Throughout surgery, mice were placed under a heating pad to avoid hypothermia. Mice were placed in a Kopf stereotaxic frame apparatus with head position to obtain a flat skull between bregma and lambda (<0.03 mm). A craniotomy was performed using a micro-drill to expose brain tissue. Dextran was delivered intra-IPN (from bregma: AP −3.45 mm; ML 0 mm; DV −5.0 mm, at a 0° angle) via a 31 G stainless steel hypodermic tube (Coopers Needleworks Ltd) connected to a Hamilton syringe controlled by an automated pump (model 220, Uno B.V, Zevenaar, NL) at an injection rate of 0.1 μl/min. Dextran delivery was followed by a 10 min diffusion period. Mice emerged from anesthesia after i.p. injection of atipamezol (Alzane, Syva, 50 mg/kg). All mice received peri- and post-operative care consisting of i.p. injection of carpofen (Carpofelican, Dechra, 5 mg/kg) on the day of surgery, followed by a maximum of 4 days of carpofen (25 mg/l) in the drinking water.

#### Immunohistochemistry

Embryos were harvested and their brains placed in ice-cold 1x PBS. Pups were decapitated and their brains placed in ice-cold 1x PBS. Adult mice were anaesthetized by i.p. injection of sodium pentobarbital (Euthanimal) and subjected to transcardial perfusion with 1x PBS followed by 4% PFA in 1x PBS, pH 7.4. Brains were fixed in 4% PFA overnight at 4°C. Subsequently, brains were placed for 48 hr in 30% sucrose for cryoprotection, frozen in - 20°C isopentane and stored at −80°C. Immunohistochemistry was performed as described previously (Brignani et al., 2020). In brief, 25 μm thick cryosections were generated and incubated in blocking solution (4% bovine serum albumin (Sigma Aldrich #05470) and 0.1% Triton x-100 (Merck Millipore #108643)) for 1 hr at RT. Sections were incubated overnight at 4°C with primary antibody diluted in blocking solution. Following washes in PBS, sections were incubated with secondary antibody in blocking solution for 1.5 hr at RT. Primary antibodies used were goat anti-CHAT (1:100, EMD Millipore, Cat#144P) and goat anti-NRP2 (1:500, R&D Systems, Cat#AF567). Secondary antibody used was donkey anti-goat Alexa Fluor 488 (1:750, Invitrogen, Cat#A11055). Sections were mounted with Fluorsave (VWR international, Cat# 345789-20). Images were taken using an Axioscope A1 microscope (Zeiss) and processed in Fiji (Schindelin et al., 2012) (version 2.0.0-rc-69/1.52o).

Experimental animals used for anatomical studies were sacrificed 48 hr to 4 days post-injection to allow for sufficient retrograde labeling of distant regions. Mice were anaesthetized by i.p. injection of sodium pentobarbital (Euthanimal, 0.1 ml), followed by transcardial perfusion with PBS and 4% PFA in PBS. Brains were dissected and post-fixed in 4% PFA overnight at 4°C. Subsequently, 30 μm sections were generated using a vibratome. Sections were collected in 1 x PBS and stained with DAPI. Sections were mounted in Fluorsave (VWR international, Cat# 345789-20) and Z-stack images were taken using a Zeiss LSM 880 laser-scanning confocal microscope.

#### BrdU labeling

Bromodeoxyuridine (BrdU) (Sigma, Cat#B5002) was diluted to 15 mg/ml in saline at 37°C for 2 hr. Pregnant C57BL/6j mice were injected i.p. with 50 mg BrdU solution per kilogram bodyweight. 8 hr to 5 days post-injection embryos were harvested and placed in ice-cold PBS. Antigen retrieval was performed by incubating the sections in 1 M HCl for 1 hr at 37°C followed by incubation in 0.1 M boric acid, pH 8.5, for 10 min at RT. ISH and IHC was performed as described above. Rat anti-BrdU (Abcam, Cat#AB6326, 1:500) and Alexa Fluor 568 donkey anti-rat (Abcam, Cat#ab175475, 1:750) antibodies were used. Sections were obtained using an Axioscope A1 microscope (Zeiss) and images were processed in Fiji (version 2.0.0-rc-69/1.52o).

#### Neurosphere assay

Cells were isolated as described previously (Guo et al., 2012b). Adult mice were sacrificed by cervical dislocation. The brains were isolated and placed in ice-cold 1 x PBS. The Hb and the subventricular zone (SVZ) were dissected out in ice cold L15++ medium (L15 (Gibco # 11415049) supplemented with 7.5 mM HEPES (Gibco #15630106) and 1 x penicillin/streptomycin (Gibco, Cat#15140148)). Tissue was dissociated using the MACS Neural Dissociation kit (Miltenyi Biotec, Cat#130-093-231) according to manufacturer’s protocol. In short, tissue was incubated in preheated dissociation solution (200 μl Trypsin and 1750 μl solution X) at 37°C for 15 minutes. 20 μl solution Y and 10 μl solution A were added per sample. Tissue was dissociated with a wide-tipped glass pipet. Samples were incubated at 37 °C for 10 minutes. The tissue was further dissociated with a series of 2 glass pipettes with descending tip sizes. 5 ml glutaMAX DMEM/F-12 (Gibco, Cat#10565018) supplemented with 1 x pen/strep was added to each sample. The suspension was filtered in a 40 μm cell strainer into a 15 ml Falcon tube and centrifuged at 300 g for 5 minutes. The cell pellet was washed with L15++ medium and centrifuged at 300 g for 5 minutes. Cells were resuspended in glutaMAX DMEM/F-12 (Gibco, Cat#10565018) supplemented with 20 ng/ml EGF (Invitrogen, Cat#PHG0311L), 20□ng/ml FGF (Gibco, Cat#PGH0367), B-27 (Gibco, Cat#17504044) and penicillin/streptomycin (Gibco, Cat#15140148) and kept in a 5% CO_2_ incubator at 37°C.

#### Brain slice preparation for electrophysiological recordings

Male and female *Cart-IRES2-cre-D^+/^:Ai14^+/-^* mice (8-12 weeks old) were anaesthetized by i.p. injection of sodium pentobarbital (Euthanimal, 0.1 ml), followed by transcardial perfusion with ice-cold, carbogen-saturated (95% O_2_ and 5% CO_2_) slicing solution containing (in mM): 92 choline chloride, 2.5 KCl, 1.2 NaH_2_PO_4_, 25 NaHCO_3_, 20 HEPES, 25 glucose, 20 N-methyl-D-glucamine (NMDG), 10 Na-ascorbate, 2 thiourea, 3 Na-pyruvate, 3.1 N-Acetyl-L-cysteine, 7 MgCl_2_.6H_2_O and 0.5 CaCl_2_.2H_2_O (mOsm 300-310; pH 7.3–7.4). After perfusion, mice were decapitated and brains removed rapidly and sectioned in ice-cold carbogen-saturated slicing solution using a vibratome (Leica VT1200S, Leica Microsystems). Coronal slices (thickness 250 μm) containing the Hb were initially recovered in carbogenated slicing solution for 5 min at 37 °C and then transferred into a holding chamber with carbogen-saturated artificial cerebrospinal fluid (ACSF), containing (in mM): 92 NaCl, 2.5 KCl, 1.2 NaH_2_PO_4_, 30 NaHCO_3_, 20 HEPES, 25 glucose, 3 Na-ascorbate, 2 thiourea, 3 Na-pyruvate, 2 MgCl_2_.6H_2_O and 2 CaCl_2_.2H_2_O (mOsm 300-310; pH 7.3–7.4). Slices were kept at RT for at least 1 hr before electrophysiological recordings. Subsequently, slices were transferred to a recording chamber, mounted on a fixed-stage, upright microscope (Olympus BX61W1), mechanically stabilized and continuously superfused (2 ml / min) with carbogenated recording ACSF containing (in mM): 124 NaCl, 2.5 KCl, 1 NaH_2_PO_4_, 26 NaHCO_3_, 5 HEPES, 11 glucose, 1.3 MgCl_2_.6H_2_O and 2.5 CaCl_2_.2H_2_O (mOsm 300-310, pH 7.3–7.4). Solution was run through an in-line heater (model SH-27B, connected to a TC-324B controller, Warner Instruments, Hamden, Connecticut, United States)) to perform electrophysiological recordings at 28–30°C.

#### Electrophysiological recordings

The Hb region was identified in brain slices under a 4x objective. Medial Hb *Cartpt^+^* cells were identified by tdTomato fluorescence and cells were visualized using infrared video microscopy and differential interference contrast (DIC) optics, under a 40x, 0.8 W water-immersion objective. Patch electrodes were pulled from borosilicate glass capillaries (Warner Instruments, Cat#300053), and had a resistance of 3–5 MΩ when back-filled with intracellular solution. A K-based internal solution was used containing (in mM): 139 K-gluconate, 5 KCl, 2 MgCl_2_. 0.2 EGTA, 10 HEPES, 10 Na creatine phosphate, 4 Mg ATP and 0.3 Na GTP (mOsm 300-310, pH 7.3-7.4, liquid junction potential of ~-12 mV). Signals were low pass filtered with a 2.9 kHz 4-pole Bessel filter and digitized at 20 kHz using an EPC-10 patch-clamp amplifier (HEKA Elektronik GmbH, Lambrecht, Rhineland-Palatinate, Germany) and PatchMaster v2.73 software.

Conventional whole-cell patch-clamp recordings were made on *Cartpt^+^* and *Cartpt^-^* neurons of the MHb ad LHb. After tight-seal (>1GΩ) cell-attached configuration, negative pressure pulses were applied to break into the cellular membrane and go to whole-cell mode. Neurons were first held in voltage clamp at −60 mV for 5 min before the start of experiments. Series resistance was not compensated and was calculated from the capacitive current amplitudes evoked by a 4 mV hyperpolarizing step and neurons that demonstrated stable holding current and series resistance were selected for further analysis. Subsequently, recordings were switched to current clamp mode. To assess membrane properties and excitability of the different neuronal populations, an IV relation was determined for all cells. Neurons were made to step from a baseline of 0 pA current injection of 400 ms to an 800 ms current injection step of −100 pA, which was followed by a return to 0 pA for 1000 ms. Every subsequent sweep had a current injection step that was 25 pA more depolarized than the previous one. Sweeps were delivered with a 10 second inter-sweep interval. Using data obtained with the IV relation, parameters of interest were calculated. Membrane resistance was calculated according to Ohm’s law, using the measured delta of steady-state voltage deflections caused by currents that did not induce active membrane conductance. Voltage sag was calculated as the delta between the minimal membrane potential and the subsequently obtained steady-state membrane potential during the 800 ms −100 pA current step. To determine action potential threshold the first IV sweep in which an action potential was triggered during the current injection step phase of 800 ms was identified and differentiated. In the differentiated waveform, the first +20 mV/ms point was identified and this timepoint was related back to the undifferentiated waveform to obtain the associated membrane potential, as the action potential threshold for the cell. Cellular capacitance was obtained by, in voltage clamp, dividing the membrane time constant tau (obtained by exponential curve fitting of the current recovery after the hyperpolarizing 4 mV test pulse towards steady-state levels) by the series resistance (obtained by dividing hyperpolarizing test pulse voltage by the amplitude of the elicited capacitive current).

### QUANTIFICATION AND STATISTICAL ANALYSIS

#### General statistical analysis

All statistical details of tests can be found in the text or figure legends. A test was considered significant when P ≤ 0.05 unless specified otherwise.

#### External datasets

For comparison we used multiple previously published datasets. Details are listed below.

##### Adult mouse Hb; Hashikawa et al 2020

For the Hashikawa *et al* dataset (GEO: GSE137478), a normalized and log-transformed Seurat object that includes the metadata on the cell clustering of the Hb cells, as described in the publication, was imported. The dataset was sequenced on the 10xGenomics platform and consisted of 5,558 cells and 19,726 genes, with a median read depth of 3,666.5 reads/cell.

##### Adult mouse Hb; Wallace et al 2020

A Seurat object was created with the Wallace *et al* dataset (Wallace et al., 2020) using the published normalized and log-transformed data matrix (https://doi.org/10.7910/DVN/2VFWF6) and the supplied metadata on the medial and lateral Hb cells. The cells were sequenced on the 10xGenomics platform. The dataset contained 3,379 cells and 25,289 genes, with a median read depth of 4,622.0 reads/cell.

##### Larval and adult zebrafish Hb; Pandey et al 2018

Zebrafish Hb datasets were used for inter-species analysis, a larval and adult zebrafish dataset from Pandey *et al* dataset (Pandey et al., 2018). Both datasets were imported by creating a Seurat object with the published normalized and log-transformed data matrix and the supplied metadata of each of the cells. The larval dataset (GEO: GSM2818523) was sequenced on the 10xGenomics platform and SmartSeq2 platform and consisted of 4,233 cells (from 10xGenomics) and 13,160 genes, with a median read depth of 2,565.94 reads/cell. The adult dataset (GEO: GSM2818522) was sequenced on the 10xGenomics platform and consisted of 7,711 cells and 9,582 genes, with a median read depth of 1,596 reads/cell.

##### Adult mouse brain tissue; Tabula Muris

As an extra control, a scRNAseq dataset from the Tabula Muris Consortium was tested (Schaum et al., 2018). The consortium performed single cell transcriptomics on cells from twenty organs and tissues, creating a compendium. For this study, FACS was used to collect cells from brain tissue. The dataset was sequenced with NovaSeq and consisted of 7,189 cells and 18,454 genes, with a median read depth of 626,662.0 reads/cell.

##### Developmental mouse hypothalamus

In the original publication (GEO: GSE132355), 129,151 developmental hypothalamus cells were sequenced at eight embryonic and four postnatal timepoints using 10xGenomics (Kim et al., 2020). This dataset was sampled to resemble the clustering of the developmental Hb dataset: fourteen clusters with a similar timepoint consistency. Additionally, 3.5% of the cells in each of those clusters were randomly sampled for the MAGMA analysis. The final object consisted of 3,385 cells and 22,187 genes, with a median read depth of 1,454.1 reads/cell.

##### Developing and adult mouse midbrain

This dataset (GEO: GSE76381) was consisted of six embryonic and one adult timepoint and used Illumina HiSeq (La Manno et al., 2016). It included in total 31 neuronal and nonneuronal cell types. The dataset consisted of 1,761 cells and 24,378 genes, with a median read depth of 8,846.0 reads/cell.

#### Mapping of scRNAseq data

Mapping was performed with the MapAndGo2 STARmap pipeline (Alemany, 2019). In short, the lanes of the fastq files were merged, then reads were demultiplexed removing all reads without a CEL-Seq2 barcode. Illumina adaptor sequences and bad quality base calls from the 3’-end were trimmed off the reads. Then reads were aligned to the reference mouse genome (Ensemble GRC38 release 93) using STAR v-2.5.3a (Dobin et al., 2013; Yates et al., 2020). Then 3 count files were produced: the unspliced file contained reads mapped to intronic regions, the spliced file contained reads mapped to exons and the total file contained both. These three files were the input for the SCANPY pipeline.

#### scRNAseq data filtering and normalization

Data were analyzed in Python (version 3.7.6) using the SCANPY package (version 1.4.6.) (Wolf et al., 2018). The *Brn3a-tLacZ* developmental dataset included 5,756 cells with a total expression of 38,068 genes, from which 85% of the reads were mapped to exonic- and 15% to intronic regions. The cells were sequenced to a median read depth of 8,482.80 reads/cell and a median of 3,156.0 genes/cell. To determine sequencing quality, External RNA Control Consortium (ERCC) spike-ins were added to each well (cell) (Jiang et al., 2011). Spike-ins are synthetic RNA with a poly-A tail, which allows their sequencing. By assessing the number of ERCC reads per cell, the sequencing quality of a cell can be determined. The dataset had a median sequencing quality of 14.10% ERCC spike-ins/cell. For further processing, cells were filtered and all the ERCC genes, together with mitochondrial genes, were removed. Cells that expressed 2,000-50,000 genes (of which less than: 15% were mitochondrial genes, 20% were ribosomal genes and 30% were ERCC spike-ins, and where at least 85% of genes were mapped to exonic regions) were kept. Besides cell filtering, we also filtered for genes that yielded the highest fraction of counts in each single cell, and genes had to be expressed in more than three cells. After filtering, the dataset contained 3304 cells with a total expression of 30,452 genes with a median read depth of 14,396.69 reads/cell, a median of 4,955.0 genes/cell and a median of 8.45% ERCC spike-ins/cell. The raw matrix with the gene expression was normalized on the total number of transcripts detected multiplied by a scaling factor of 10,000. The data were then log transformed (ln) after the addition of a pseudo-count of 1. Additionally, cells that did not express *Pou4f1/Brn3a* (0.0 > *Pou4f1*) were removed. These uncorrected data were stored as a raw count matrix which was used to calculate DEGs, plot gene expression levels and perform downstream analysis.

The E18 WT dataset contained 1920 cells and 36,196 genes. 85% of the reads were spliced and 15% of the reads were unspliced. The median number of genes per cell was 2857. We filtered out ERCC genes, mitochondrial genes, genes that were detected in less than 3 cells and cells that expressed less that 2000 genes. After filtering, the dataset contained 1002 cells and 28237 genes. Next, we removed cells that did not meet the following criteria: number of counts < 50000, mitochondrial reads < 15%, ribosomal reads < 20%, ERCC genes < 30% and coding reads > 85%. After filtering the dataset contained 948 cells and 28237 genes. The data were normalized to 10,000 reads per cell and log-transformed. Genes with an expression between 0.0125 and 3 and a minimum dispersion of 0.5 were considered highly variable and were kept in the dataset for clustering. Then the effects of total counts per cell and percentage mitochondrial reads were regressed out and data were scaled to a max value of 10.

#### Batch effect correction for the devHb dataset using Seurat v3

Seurat v3 integration software (Stuart et al., 2019a, b) requires a Seurat object that is normalized and log-transformed (raw matrix) as input. The CCA, L2-normalization, finding the MNNs and scoring them was executed using the FindTransferAnchors function, where thirty, fifteen and four MNNs were used for filtering-, scoring- and finding anchors, respectively. The IntegrateData function merged the data and corrected the data matrix, which was then stored in the integrated assay in the Seurat object. The batch-corrected count matrix was then imported into SCANPY where it replaced the uncorrected count matrix. This matrix was used for downstream clustering.

#### Principal Component Analysis and clustering

A Principal Component Analysis (PCA) was computed with fifty components. Then the nearest neighbors were calculated using the first forty principal components. Subsequently, t-SNE coordinates were computed using the first forty principal components. Clustering was performed with the Louvain algorithm (Traag, 2017).

#### Differential gene expression

Marker genes per Louvain cluster were calculated with the Wilcoxon-rank-sum test using the sc.tl.rank_genes_groups() function in SCANPY. In order to find more specific marker genes, as depicted in Fig. 1F and S8C, markers genes were filtered using the following parameters: min_in_group_fraction = 0.5, max_out_group_fraction = 0.3, min_fold_change = 1.5. Adjusted p-values of DEGs between the left and right Hb were derived using the sc.get.rank_genes_groeps_df() function.

#### scVELO

Filtered and normalized E18 data were used as a input for the scVELO (Bergen et al., 2020) implementation in SCANPY. First- and second-order moments were calculated using the first 30 principle components and 30 neighbors. Velocity was computed using the “stochastic” mode.

#### CellRank trajectory inference

Cellrank trajectory inference was performed on the developmental dataset(Lange et al., 2020). First, it computed terminal states, the number of terminal states was defined by the user (n = 7) using the cr.tl.terminal_states() function. Then, it defined initial states with the cr.tl.initial_states() function. Next, fate maps were computed to define the likelihood that an individual cell reaches a certain fate. Subsequently, the individual fate maps were aggregated to cluster-level fate maps by calculating latent time and running PAGA. Gene expression trends along pseudo- or latent time were calculated based on a general additive model (GAM) with the cr.pl.gene_trends() function.

#### Correlation analysis

In a correlation analysis, the gene expression patterns of clusters from two datasets (referred to as dataset 1 and 2) were correlated. This analysis was based on previously described methods (Tosches et al., 2018). DEG were used for the calculation of pairwise cluster correlations because those suffice as cell type markers, possibly even cross-species. DEG were computed with the MAST (Finak et al., 2015) package using the zlm() function. This function uses count data to model the expression based on cluster membership and number of detected genes per cell. First, DEG were computed within each dataset, by calculating among all possible pairs of clusters. Genes were only selected if they were detected in more than 40% of cells, in at least one of the two compared clusters and if there was a minimum average difference of ln(2) between the two clusters, together with a false discovery rate (FDR) of 5%. As input, the function requires data formatted as a SingleCellExperiment (Amezquita et al., 2020) such as a Seurat object. For the developmental Hb data, the SCANPY object was made into a Seurat object by exporting the expression matrix, the raw matrix and the metadata. Together, these were transformed into a Seurat object using the function CreateSeuratObject(), where the batch-corrected matrix was added separately. After DEG were determined for each dataset, the intersection of DEG sets for dataset 1 and 2 was taken as input for the correlation analysis. By doing so, genes absent in either one of the datasets were excluded from the analysis. Gene names in the zebrafish datasets were converted to one-to-one mouse orthologous gene using BioMarts (Kinsella et al., 2011). With the AverageExpression() function in Seurat a data matrix was created that holds the average expression for the DEGs of each cluster. To show the relationship of the genes to each of the clusters, a gene-cell type specificity matrix was calculated. Each average expression was divided by the sum of all, giving a fraction of specificity for all clusters.

Pairwise Spearman rank order correlations were calculated on the gene-cell type specificity matrices, between all clusters for dataset 1 and 2. Significance of correlation coefficients was determined with a permutation test, where the gene expression values were shuffled one thousand times across cell types. Afterwards, the resulting Spearman correlation coefficient rho was calculated with the function:

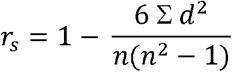

Where r_s_, d and n resemble rho, the difference between ranks and number of genes, respectively. The *P* value was calculated as the fraction of the absolute value of rho values that were greater than or equal to the value of rho for the non-shuffled data. The correlation coefficients and *P* values were plotted in a heatmap from the heatmaply^55^ package, where the clusters were ordered by hierarchical clustering, with Euclidean metric for distance calculation and complete-linkage clustering.

#### MAGMA gene property analysis

Genome Wide Association Studies (GWAS) summary statistics on MDD (Wray et al, 2018), SCZ (Ripke et al., 2014), BMI (Locke et al., 2015), BIP (Hou et al., 2016) and height (Wood et al., 2014) were analyzed in MAGMA (de Leeuw et al., 2015; Watanabe et al., 2019) to converge genetic associations to cell types.

The GWAS datasets had to be processed into the same format including the SNP ID, chromosome and base pair number as the first three columns, and the *P* value (decimal notation) as column “P”. A gene-cell type specificity matrix, holding the relationship of the genes to each of the clusters, was calculated. After the matrix was created, uninformative genes were removed from the matrix using an ANOVA test of variance. This analysis was limited to one-to-one human orthologous genes to match GWAS loci. This resulted in the following number of genes for each dataset matrix: 13,330 (DevHb), 11,392 (Hashikawa et al., 2020), 11,217 (Wallace et al., 2020), 14,033 (Schaum et al., 2018), 11,381 (Kim et al., 2020) and 12,214 (La Manno et al., 2016). Gene-property analysis was performed for each GWAS and differently processed scRNAseq dataset combination, where the input of the function was given as follows: --gene-results [GWAS] --gene-covar [scRNAseq]. The direction of gene property testing was set to positive (using the flag --model direction=greater). A positive gene-property analysis was performed. The results of the gene-property analysis were visualized in a bar plot, facet wrapped on GWAS type, using ggplot2 (Dandekar et al., 2009). The significance threshold was corrected using a Bonferroni correction.

#### Quantification of cell body size

To quantify cell-body size (Fig. 7C), 2x digital zoom DIC images of *Cartpt^+^* and *Cartpt^-^* neurons were processed in ImageJ. Ferret’s diameter (the longest distance between any two points) was measured along each cell-containing ROIs. One-way ANOVA was performed comparing *Cartpt^+^* neurons (n = 15 cells, N = 8 mice), *Cartpt^-^* MHb neurons (n = 14 cells, N = 8 mice) and *Cartpt^-^* LHb neurons (n = 5 cells, N = 2 mice). Results are shown as mean ± SEM.

#### Two-step cluster analysis of electrophysiological parameters

Two-step cluster analysis (Fig. 7H-J) was based on the Schwarz’s Bayesian criterion, with automatic generation of cluster numbers to avoid biased cluster-member selection. Cell capacitance and membrane resistance were chosen as input variables for clustering, as these parameters showed the largest differences between *Cartpt^+^* neurons and both *non-Cartpt* populations. Statistical analysis, including cluster analysis, was performed using IBM SPSS Statistics (IBM Corp., Armonk, NY, US, version 27) software.

#### Quantification of Dextran^+^/Cartpt^+^ neurons

Corresponding to Fig. 6F. For quantification of Dextran^+^/*Cartpt^+^* cells, representative samples (n = 2 mice, N = 19 slices) containing the border region between the MHb and LHb were imaged and analyzed by an automated in-house Fiji (Schindelin et al., 2012) script that allowed for manual inspection and exclusion of false-positive cell identification. In brief, using 40x maximum projection images, Dextran^+^ cells (n = 175) were identified by sequential application of “Intermodes” and “Max entropy” auto-thresholds, with inclusion parameters of particles size (60–6000) and circularity (0–1.00). ROI masks of detected Dextran^+^ cells were then projected to the TdTomato (*Cartpt^+^*) channel and overlapping signals were established in an unbiased manner. No further statistical tests were applied to these data.

#### 3DISCO analysis of Cartpt^+^ neuron localization

Corresponding to Fig. 6E. Z-stack light sheet images were merged into an 3D Imaris file. Then, surfaces were created of the 3^rd^ ventricle and the left half of the Hb. This was done in the “surfaces” tool in Imaris, using the “manual creation” option, with “drawing mode = click” in the “surfaces” window. The surfaces were traced in coronal planes every 100 μm along the rostral-to-caudal axis. Then, the *Cartpt^+^* RFP^+^ channel was masked using the Hb surface, creating a new masked *Cartpt^+^* RFP^+^ channel. Subsequently, Cartpt^+^ RFP^+^ neurons were marked with spots. The average diameter of the *Cartpt^+^* cells was measured in the “slice view” setting. This average diameter was used as input for creating the spots. Spots were created from the *Cartpt^+^* RFP^+^ source channel with background subtraction. Then, spots were filtered for the mean intensity in the masked *Cartpt^+^* RFP^+^ channel. As a final check, spots were manually inspected for false positives / false negatives. To localize the *Cartpt^+^* spots, a reference frame was placed at the most rostral edge of the Hb aligned along the midline of the 3^rd^ ventricle. To measure the length of the Hb along the rostral-to-caudal axis the most caudal point on the edge of the Hb surface was marked with a spot. Then, 4 equal-sized bins were created along this axis. This was done filtering the spots based on “position Y reference frame”, adding an upper and lower boundary per bin. Spots within each bin were color-coded. Then, using the “statistics” option within the *Cartpt^+^* spots object the “distance from surface = 3^rd^ ventricle” was used to calculate the distance from the ventricle for each spot (medial-to-lateral axis X). These results were exported as *.cvs* and analyzed in Graphpad version 9.1.1. No further statistical tests were applied to these data.

#### Statistical analysis electrophysiological parameters

Electrophysiological parameters were calculated off-line using Igor Pro 8 (version 8.0.4.2, Wavemetrics, Tigard, OR, US). Statistical tests on these electrophysiological parameters were subsequently performed in IBM SPSS 27. Results were shown as mean ± SEM. Per experiment statistical “n” and test were indicated in the text or legend.

