## Supplementary figures for "Molecular Signatures and Cellular Diversity During Mouse Habenula Development"

Figure S1

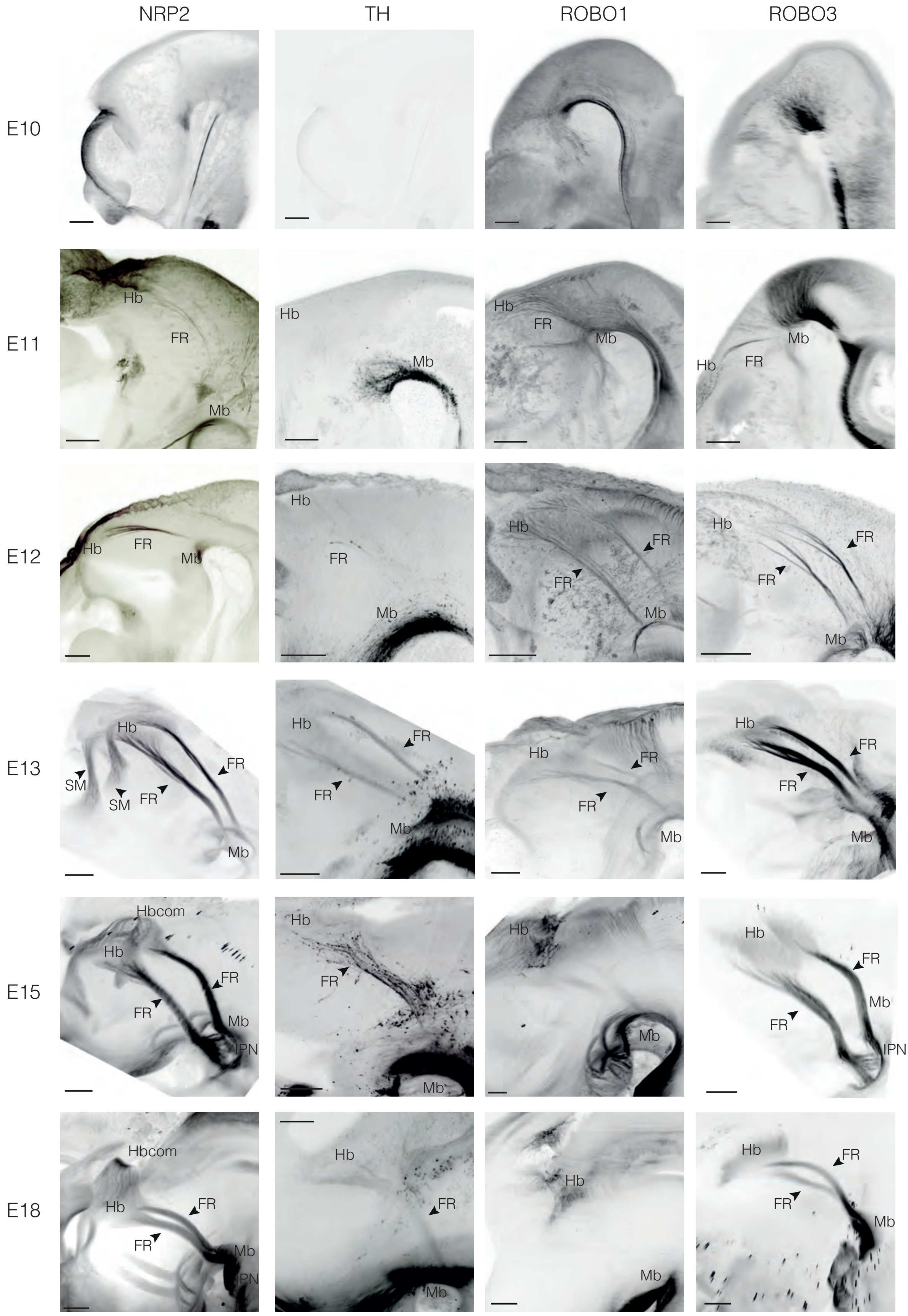

**Supplementary Figure S1. Three-dimensional visualization of important hallmarks of mouse habenula development. Related to Figure 1.** Whole-mount immunostaining for neuropilin-2 (NRP2), tyrosine hydroxylase (TH), roundabout1 (ROBO1) or ROBO3 of E10, E11, E12, E13, E15 and E18 mouse embryos followed by 3DISCO tissue clearing and fluorescent light sheet microscopy (FLSM). Rostral is to the left and dorsal to the top. Scale bar, 200 µm. FR, fasciculus retroflexus; Hb, habenula; Hbcom, habenula commissure; IPN, interpeduncular nucleus; Mb, midbrain; SM, stria medularis.

Figure S2

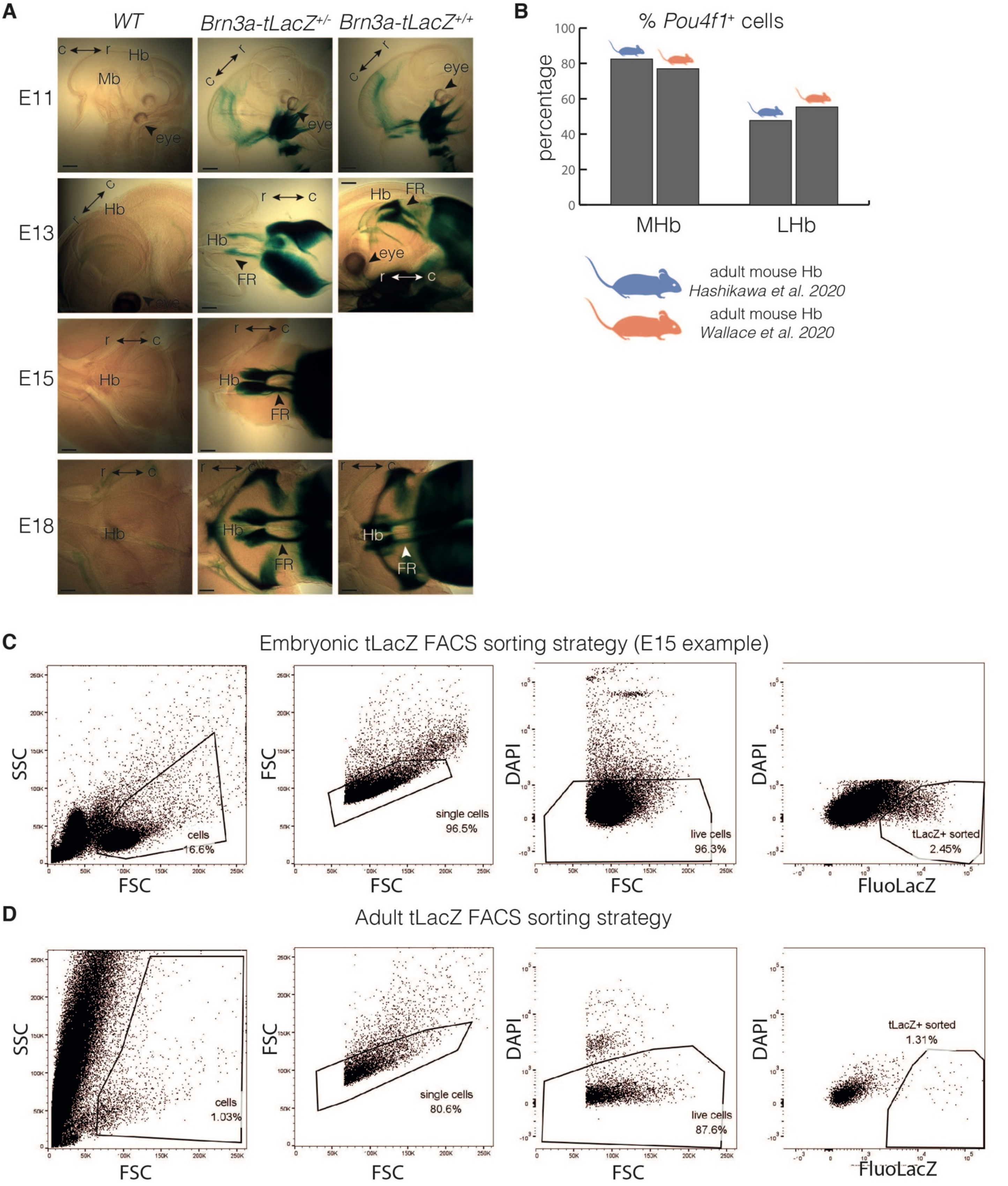

**Supplementary Figure S2. FACS-based strategy to isolate post-mitotic habenula neurons from *Brn3a-tauLacZ mice*. Related to Figure 1.**

**A,** Brn3a$^+$ neurons and axonal projections are labeled throughout embryonic development in *Brn3a-tLacZ* mice. β-gal staining of cleared *Brn3a-tLacZ*$^{+/-}$ and *Brn3a-tLacZ*$^{-/-}$ mice shows β-gal expression in *Brn3a-tLacZ*$^{+/-}$ but not wild-type (WT; *Brn3a-tLacZ*$^{-/-}$) embryos. Rostral to caudal axis is show in sagittal plane (E11, E13) and horizontal place (E15, E18).

**B,** Percentage of *Pou4f1*$^+$ cells in the medial (MHb) and lateral habenula (LHb) in two adult mouse habenula scRNAseq datasets (Hashikawa et al., 2020; Wallace et al., 2020).

**C, D,** FACS gating strategy to obtain embryonic (C) and adult (D) Brn3a-tauLacZ$^+$ habenula neurons. Gating selects for cells, single cells, live cells and tauLacZ$^+$ cells, respectively. C, caudal; E, embryonic day; FACS, fluorescence-activated cell sorting; FSC, forward scatter; FR, fasciculus retroflexus; Hb, habenula; IPN, interpeduncular nucleus; Mb, midbrain; P2, prosomere 2; r, rostral; SM, stria medularis; SSC, side scatter. Scale bar = 200 μm.

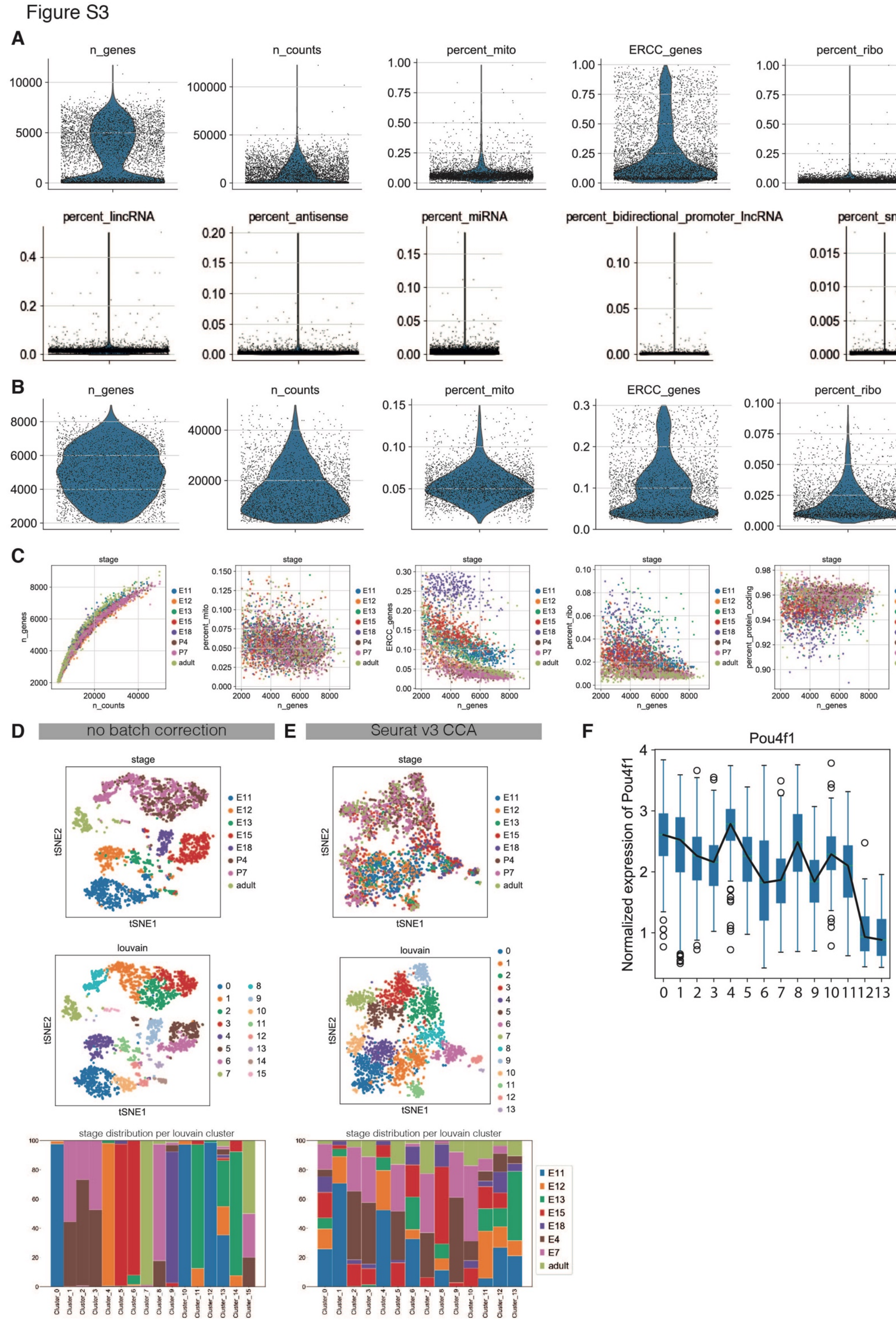

**Supplementary Figure S3. Quality control scRNA-seq data. Related to Figure 1.**

**A, B,** Violin plots showing the distribution of cells across multiple features before (A) and after (B) filtering. A relatively high number of reads is detected per cell while the percentage of mitochondrial and ribosomal reads is relatively low, indicating that the quality of the cells in good (A). After filtering for cells with more than 2000 genes, a max number of 8000 genes per cell is detected (B).
**C,** Dot plots showing the number of genes and the percentage of mitochondrial reads, ERCC reads, ribosomal reads and protein coding reads of the identified cells. Colors indicate different developmental stage. After filtering, the percentage of mitochondrial and protein coding genes is comparable across ages (B).
**D, E**, Removal of timepoint batch effects using Seurat v3 CCA. t-SNE embedding showing developmental timepoints (top) and clusters (middle) before (D) and after (E) batch effect correction. In the bottom figure, the distribution of cells from each developmental stage per cluster before or after batch effect correction is shown.
**F,** Boxplot of normalized *Pou4f1* expression levels per cluster. *Pou4f1* expression is found in all clusters. PC clusters 12 and 13 express low levels of *Pou4f1*, meaning that we probably sampled these during the process of commitment to habenula fate.
Mito, mitochondrial reads; n, number; ribo, ribosomal reads; E, embryonic day; P, postnatal day; CCA, canonical correlation analysis.

### Figure S4

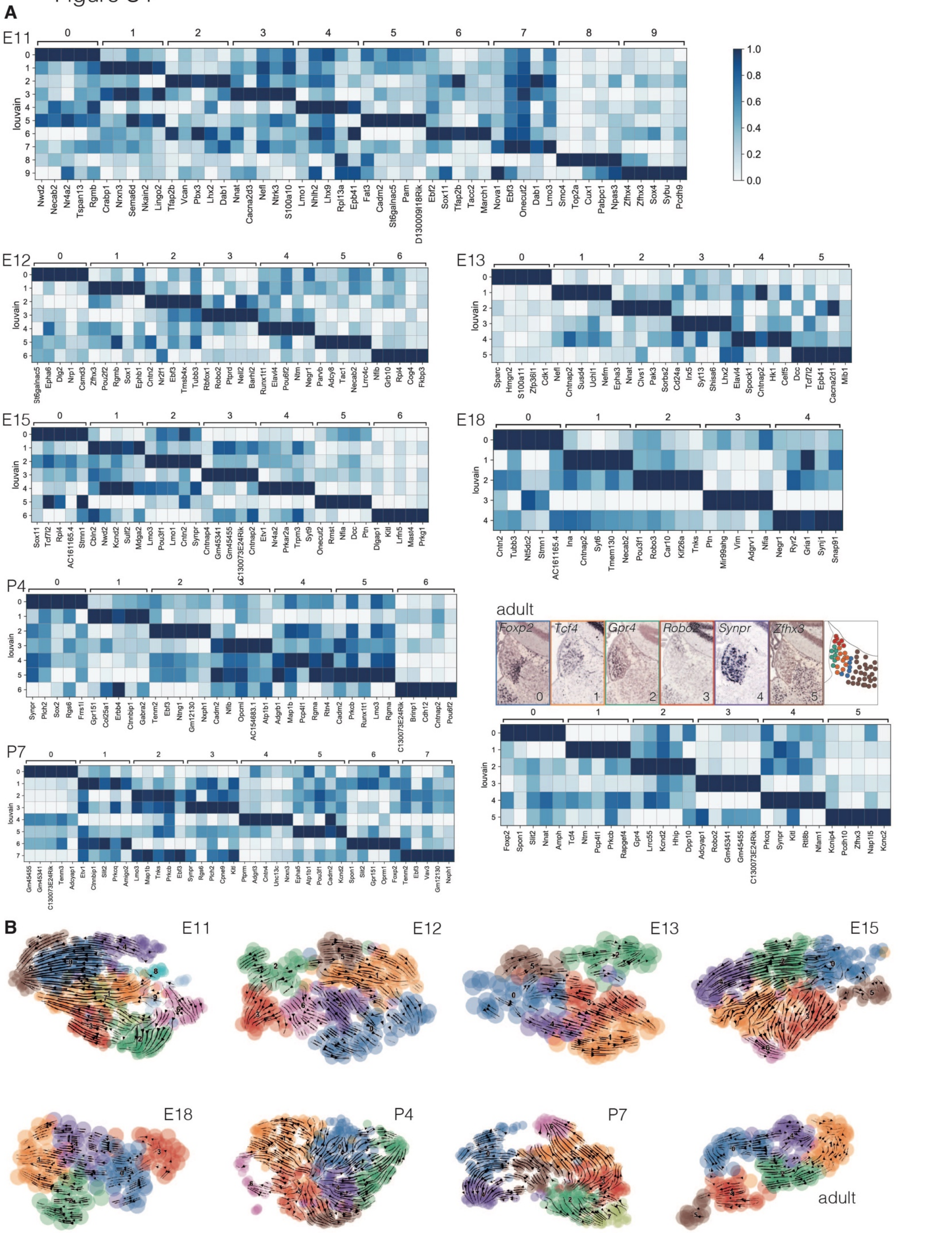

**Supplementary Figure S4. Subclustering of cells per timepoint. Related to Figure 1.**

**A,** Matrix plot of scaled expression levels of marker genes per subcluster per timepoint. For adult, *in situ* hybridization data (Allen Brain Atlas) is shown of marker genes per subcluster. *In situ* hybridization data are summarized in a color-coded graphical overview at the right. Colors indicate the adult Louvain cluster as depicted in (B). E, embryonic day; P, postnatal day.

**B,** scVELO analysis of Louvain subclusters shown in A per sampled timepoint.

Figure S5

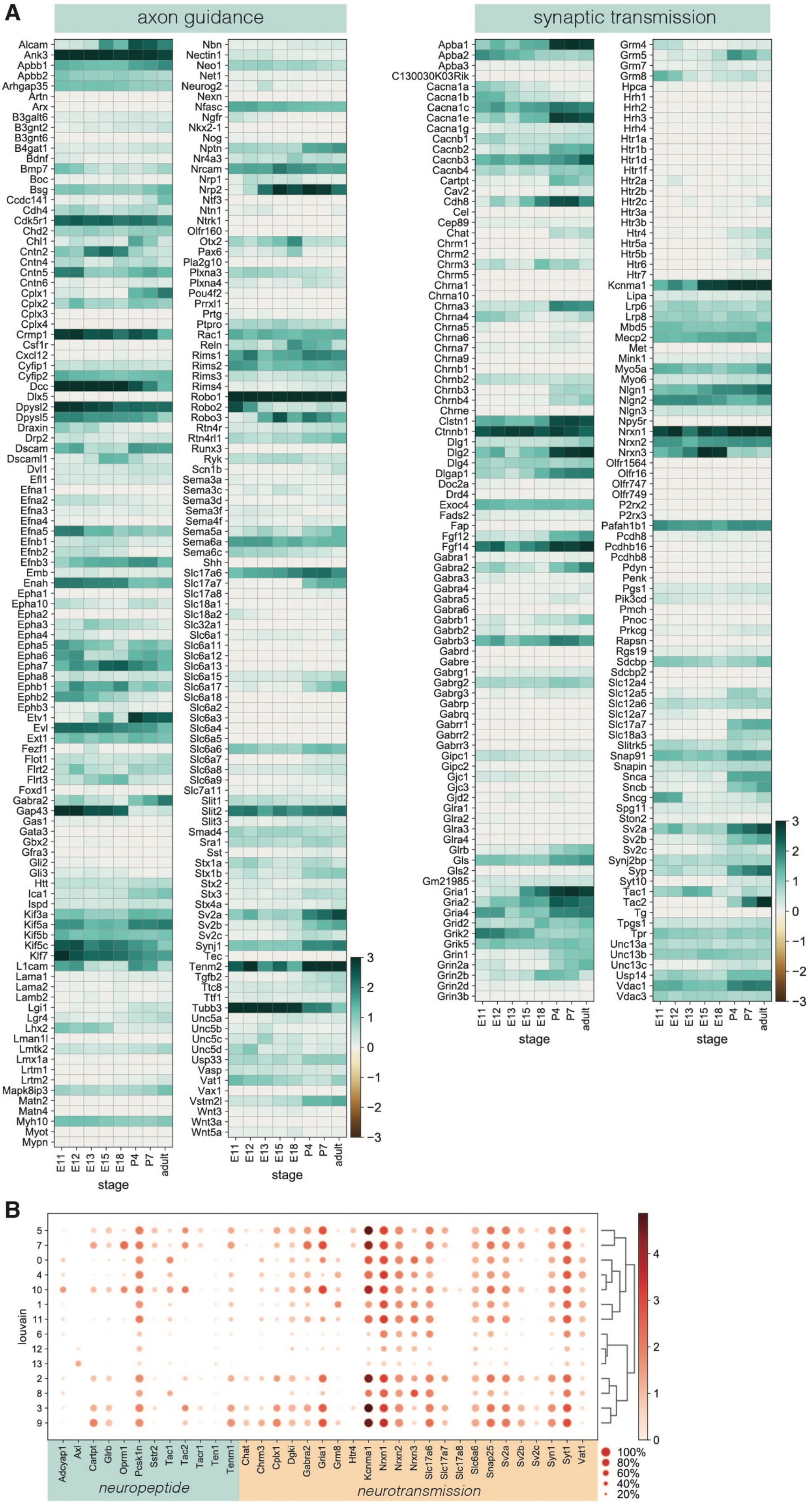
A
axon guidance
synaptic transmission
Alcam
Ank3
Apbb1
Apbb2
Arhgap35
Artn
Arx
B3galt6
B3gnt2
B3gnt6
B4gat1
Bdnf
Bmp7
Boc
Bsg
Ccdc141
Cdh4
Cdk5r1
Chd2
Chl1
Cntn2
Cntn4
Cntn5
Cntn6
Cplx1
Cplx2
Cplx3
Cplx4
Crmp1
Csf1r
Cxcl12
Cyfip1
Cyfip2
Dcc
Dlx5
Dpysl2
Dpysl5
Draxin
Drp2
Dscam
Dscaml1
Dvl1
Efl1
Efna1
Efna2
Efna3
Efna4
Efna5
Efnb1
Efnb2
Efnb3
Emb
Enah
Epha1
Epha10
Epha2
Epha3
Epha4
Epha5
Epha6
Epha7
Epha8
Ephb1
Ephb2
Ephb3
Etv1
Evl
Ext1
Fezf1
Flot1
Flrt2
Flrt3
Foxd1
Gabra2
Gap43
Gas1
Gata3
Gbx2
Gfra3
Gli2
Gli3
Htt
Ica1
Ispd
Kif3a
Kif5a
Kif5b
Kif5c
Klf7
L1cam
Lama1
Lama2
Lamb2
Lgi1
Lgr4
Lhx2
Lman1l
Lmtk2
Lmx1a
Lrtm1
Lrtm2
Mapk8ip3
Matn2
Matn4
Myh10
Myot
Mypn
Nbn
Nectin1
Neo1
Net1
Neurog2
Nexn
Nfasc
Ngfr
Nkx2-1
Nog
Nptn
Nr4a3
Nrcam
Nrp1
Nrp2
Ntf3
Ntn1
Ntrk1
Olfr160
Otx2
Pax6
Pla2g10
Plxna3
Plxna4
Pou4f2
Prrxl1
Prtg
Ptpro
Rac1
Reln
Rims1
Rims2
Rims3
Rims4
Robo1
Robo2
Robo3
Rtn4r
Rtn4rl1
Runx3
Ryk
Scn1b
Sema3a
Sema3c
Sema3d
Sema3f
Sema4f
Sema5a
Sema6a
Sema6c
Shh
Slc17a6
Slc17a7
Slc17a8
Slc18a1
Slc18a2
Slc32a1
Slc6a1
Slc6a11
Slc6a12
Slc6a13
Slc6a15
Slc6a17
Slc6a18
Slc6a2
Slc6a3
Slc6a4
Slc6a5
Slc6a6
Slc6a7
Slc6a8
Slc6a9
Slc7a11
Slit1
Slit2
Slit3
Smad4
Sra1
Sst
Stx1a
Stx1b
Stx2
Stx3
Stx4a
Sv2a
Sv2b
Sv2c
Synj1
Tec
Tenm2
Tgfb2
Ttc8
Ttf1
Tubb3
Unc5a
Unc5b
Unc5c
Unc5d
Usp33
Vasp
Vat1
Vax1
Vstm2l
Wnt3
Wnt3a
Wnt5a
Apba1
Apba2
Apba3
C130030K03Rik
Cacna1a
Cacna1b
Cacna1c
Cacna1e
Cacna1g
Cacnb1
Cacnb2
Cacnb3
Cacnb4
Cartpt
Cav2
Cdh8
Cel
Cep89
Chat
Chrm1
Chrm2
Chrm3
Chrm5
Chrna1
Chrna10
Chrna3
Chrna4
Chrna5
Chrna6
Chrna7
Chrna9
Chrnb1
Chrnb2
Chrnb3
Chrnb4
Chrne
Clstn1
Ctnnb1
Dlg1
Dlg2
Dlg4
Dlgap1
Doc2a
Drd4
Exoc4
Fads2
Fap
Fgf12
Fgf14
Gabra1
Gabra2
Gabra3
Gabra4
Gabra5
Gabra6
Gabrb1
Gabrb2
Gabrb3
Gabrd
Gabre
Gabrg1
Gabrg2
Gabrg3
Gabrp
Gabrq
Gabrr1
Gabrr2
Gabrr3
Gipc1
Gipc2
Gjc1
Gjc3
Gjd2
Glra1
Glra2
Glra3
Glra4
Glrb
Gls
Gls2
Gm21985
Gria1
Gria2
Gria4
Grid2
Grik2
Grik5
Grin1
Grin2a
Grin2b
Grin2d
Grin3b
Grm4
Grm5
Grm7
Grm8
Hpca
Hrh1
Hrh2
Hrh3
Hrh4
Htr1a
Htr1b
Htr1d
Htr1f
Htr2a
Htr2b
Htr2c
Htr3a
Htr3b
Htr4
Htr5a
Htr5b
Htr6
Htr7
Kcnma1
Lipa
Lrp6
Lrp8
Mbd5
Mecp2
Met
Mink1
Myo5a
Myo6
Nlgn1
Nlgn2
Nlgn3
Npy5r
Nrxn1
Nrxn2
Nrxn3
Olfr1564
Olfr16
Olfr747
Olfr749
P2rx2
P2rx3
Pafah1b1
Pcdh8
Pcdhb16
Pcdhb8
Pdyn
Penk
Pgs1
Pik3cd
Pmch
Pnoc
Prkcg
Rapsn
Rgs19
Sdcbp
Sdcbp2
Slc12a4
Slc12a5
Slc12a6
Slc12a7
Slc17a7
Slc18a3
Slitrk5
Snap91
Snapin
Snca
Sncb
Sncg
Spg11
Ston2
Sv2a
Sv2b
Sv2c
Synj2bp
Syp
Syt10
Tac1
Tac2
Tg
Tpgs1
Tpr
Unc13a
Unc13b
Unc13c
Usp14
Vdac1
Vdac3
E11
E12
E13
E15
E18
P4
P7
adult
stage
3
2
1
0
−1
−2
−3
B
louvain
5
7
0
4
10
1
11
6
12
13
2
8
3
9
Adcyap1
Axl
Cartpt
Glrb
Oprm1
Pcsk1n
Sstr2
Tac1
Tac2
Tacr1
Ten1
Tenm1
Chat
Chrm3
Cplx1
Dgki
Gabra2
Gria1
Grm8
Htr4
Kcnma1
Nrxn1
Nrxn2
Nrxn3
Slc17a6
Slc17a7
Slc17a8
Slc6a6
Snap25
Sv2a
Sv2b
Sv2c
Syn1
Syt1
Vat1
neuropeptide
neurotransmission
4
3
2
1
0
100%
80%
60%
40%
20%

**Supplementary Figure S5. Analysis of developmental stage- or cluster-specific gene expression related to axon guidance and synaptic transmission. Related to Figure 2.**

**A,** Heatmap showing expression levels per sampling timepoint (developmental stage) of transcripts involved in axon guidance (GO:0008046, GO:0007411) and synaptic transmission (GO:0007268).

**B,** Dot plot showing the expression levels of neuropeptide- and neurotransmission-related transcripts per Louvain cluster.

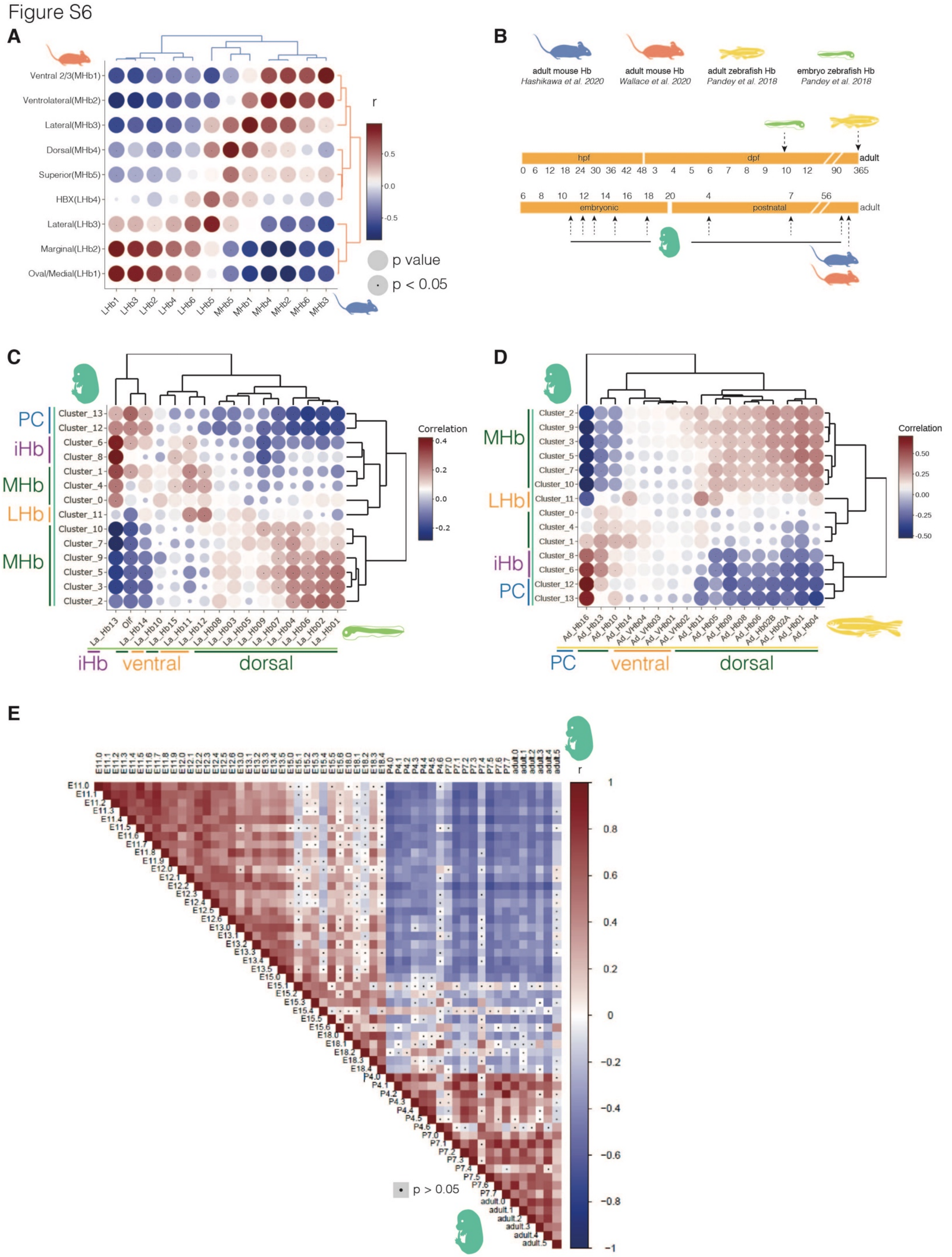

**Supplementary Figure 6. Comparison of developing mouse habenula neurons to adult mouse and zebrafish habenula neurons. Related to Figure 2.**

**A,** Matrix plot showing the correlation between clusters of two previously published adult habenula datasets: Wallace et al., 2020 (orange) and Hashikawa et al., 2020 (blue). Disk size indicates *P* value. Black dot in the disk center indicates a statistically significant correlation ($P < 0.05$). Labeling of clusters is as published in the indicated papers.
**B,** Graphical overview of mouse and zebrafish development. Arrows indicate sampling timepoints of each dataset.
**C, D,** Matrices showing the correlation between developing mouse habenula neuronal cell populations and larval zebrafish (C) and adult zebrafish (D) habenula cell populations. Disk size indicates *P* value. Black dot in the disk center indicates a statistically significant correlation ($P < 0.05$).
**E,** Matrix indicating the correlation between subclusters of developing mouse habenula populations (as defined in **Figure S4**). Black dot in the square center indicates a statistically insignificant correlation ($P > 0.05$). Ad, adult; dpf, days post-fertilization; E, embryonic day; Hb, habenula; hpf, hours post-fertilization; iHb, immature habenula; La, larval; LHb, lateral habenula; MHb, medial habenula; Olf, olfactory neurons; P, postnatal day; PC, progenitor cells.

Figure S7

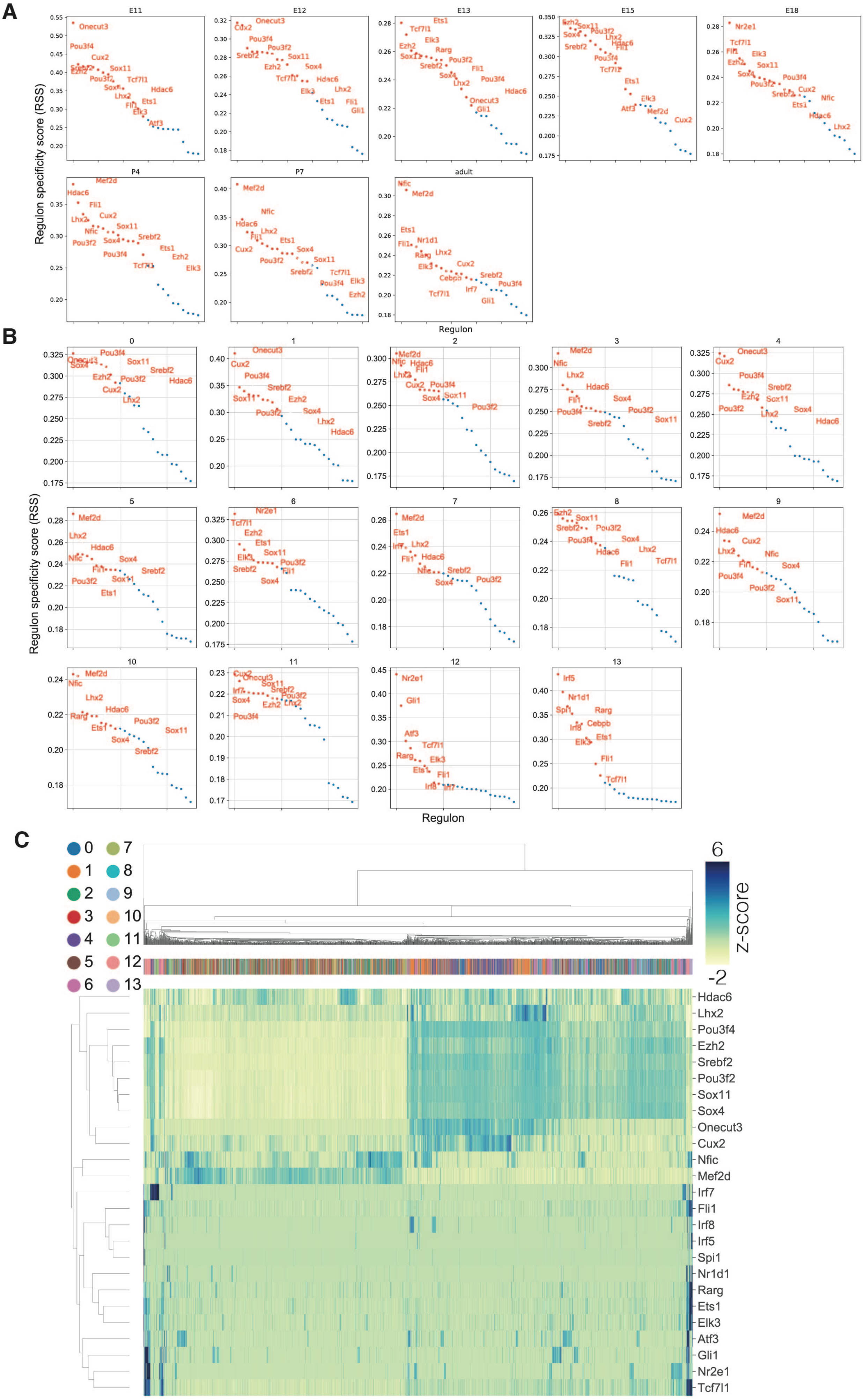
A
E11
E12
E13
E15
E18
P4
P7
adult
Regulon specificity score (RSS)
Regulon
B
0
1
2
3
4
5
6
7
8
9
10
11
12
13
Regulon specificity score (RSS)
Regulon
C
0
1
2
3
4
5
6
7
8
9
10
11
12
13
6
z-score
-2
Hdac6
Lhx2
Pou3f4
Ezh2
Srebf2
Pou3f2
Sox11
Sox4
Onecut3
Cux2
Nfic
Mef2d
Irf7
Fli1
Irf8
Irf5
Spi1
Nr1d1
Rarg
Ets1
Elk3
Atf3
Gli1
Nr2e1
Tcf7l1

**Supplementary Figure 7. pySCENIC gene regulatory network analysis of developing mouse habenula neurons. Related to Figure 4.**
**A, B,** Regulon specificity scores (RSS) per developmental stage (A) and Louvain cluster (B).
**C,** Heatmap showing the Z-score per cell per regulon. Cells are color-coded by Louvain cluster. E, embryonic day; P, postnatal day; RSS, Regulon specificity scores.

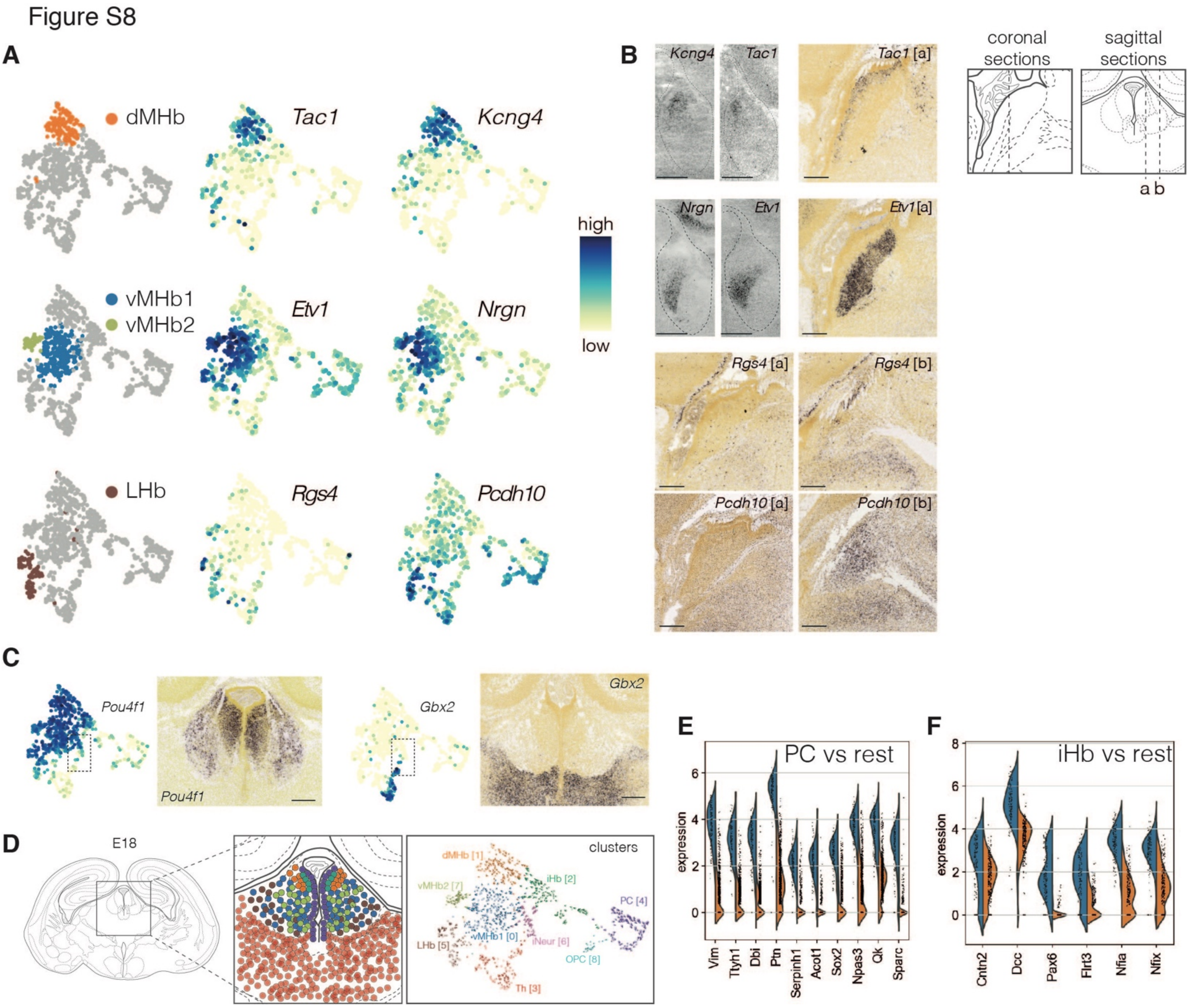

**Supplementary Figure 8. scRNAseq analysis of the whole E18 mouse habenula. Related to Figure 5.**
**A,** Left: indication of E18 WT clusters on UMAP embedding. Right: expression patterns of marker genes on UMAP embedding.
**B,** *In situ* hybridization of marker genes at E18 in sagittal sections (location [a] and [b] from the Allen Brain Atlas). Scale bar = 200 µm.
**C,** Right: expression patterns of marker genes on UMAP embedding. Left: indication of cluster on UMAP embedding.
**D,** Graphical overview of all E18 cell populations as defined by the Louvain algorithm including their location in the habenula.
**E, F,** Top 10 DEGs defining PCs (E) and top 5 DEGs defining iHb neurons (F) in comparison to all other cells in the E18 WT dataset calculated with Wilcoxon rank sum test.
Hb, habenula; iHb, immature habenula; iNeur, immature neurons; LHb, lateral habenula; MHb, medial habenula; OPC, oligodendrocyte precursor cells; PC, progenitor cells; Th, thalamus.

Figure S9

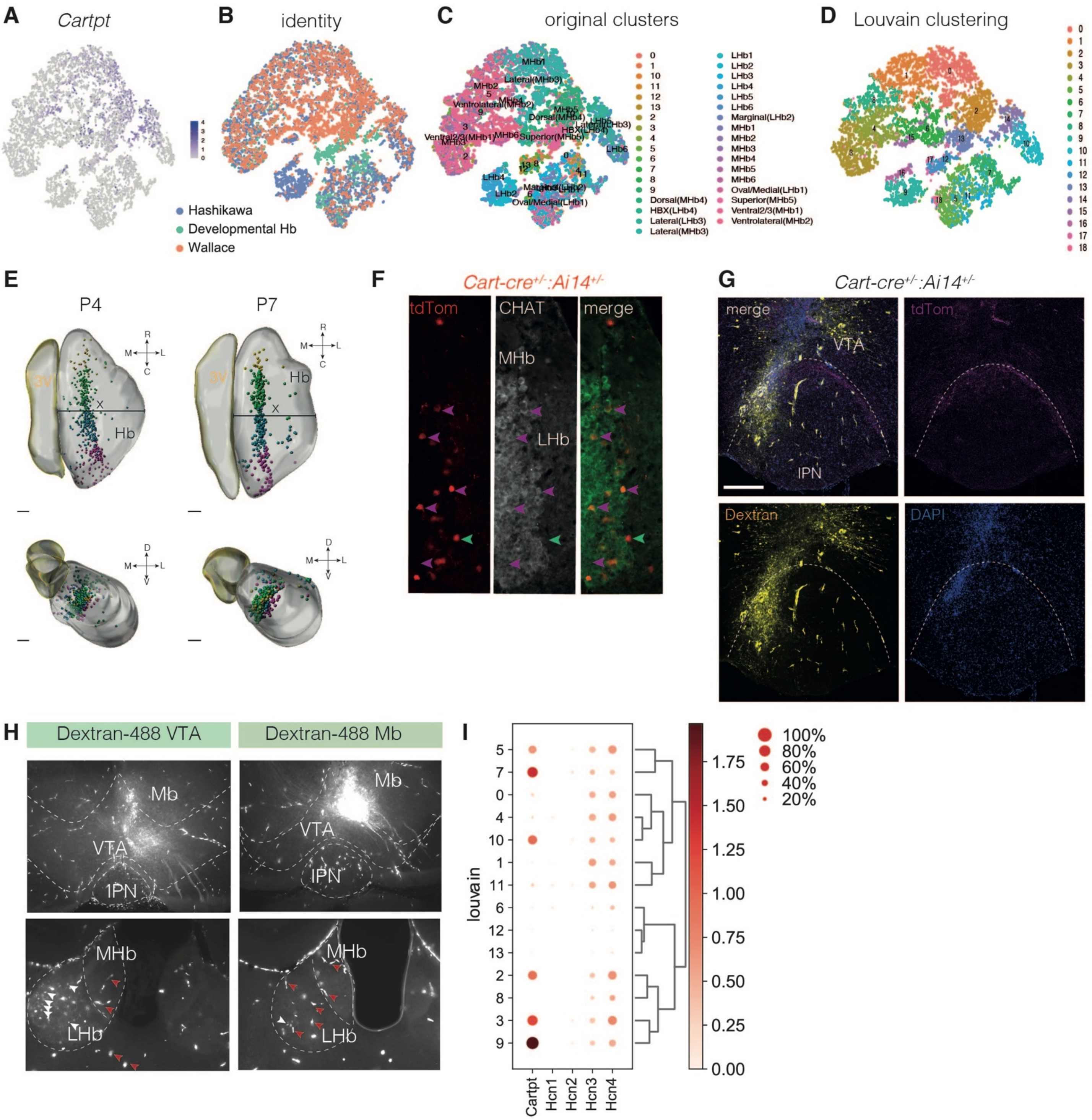

**Supplementary Figure 9. Molecular signature, projection target and electrophysiological properties of Cartpt$^+$ habenula neurons. Related to Figure 6 and 7.**

**A-D,** t-SNE embedding with merged habenula cells from adult mouse habenula (Wallace et al., 2020; Hashikawa et al., 2020) and DevHb datasets. (A) *Cartpt* expression levels. Cells are color-coded by original identity (B), original cluster (C) and new Louvain clustering (D) after merging.

**E,** Whole-mount immunostaining for tdTomato was performed in P4 and P7 *Cart-cre:Ai14* mice followed by 3DISCO tissue clearing and fluorescent light sheet microscopy (FLSM). Top: horizontal plane. Bottom: coronal plane. The third ventricle is masked in yellow and the habenula is masked in grey. For quantification of tdTomato$^+$ neuron localization the habenula is subdivided in 4 equally-sized bins. Here shown for the horizontal plane. Spots are color-coded per bin: yellow, green, blue and pink from a rostral-to-caudal perspective. Scale bar = 100 µm.

**F,** Immunohistochemistry for tdTomato and CHAT in a coronal section of adult *Cart-cre:Ai14* mice showing the border region. Purple arrows indicate tdTomato$^{+}$/CHAT$^{+}$ double positive neurons, green arrows indicate tdTomato$^{+}$/CHAT$^{-}$ single-positive neurons. Scale bar = 200 µm.

**G, H,** Dextran-488 injections in the VTA and midbrain hitting the most dorsal edge of the VTA of adult *Cart-cre:Ai14* mice (G) result in Dextran-488$^+$ neurons in the LHb. White arrowheads indicate neurons, read arrowheads indicate blood vessels.

**I,** Dotplot showing expression levels of *Cartpt* and *Hcn1-4* per Louvain cluster (DevHb dataset).

E, embryonic; FR, fasciculus retroflexus; Hb, habenula; IPN, interpeduncular nucleus; LHb, lateral habenula; Mb, midbrain; MHb, medial habenula; P, postnatal; tdTom, tdTomato; VTA, ventral tegmental area.

#### Figure S10

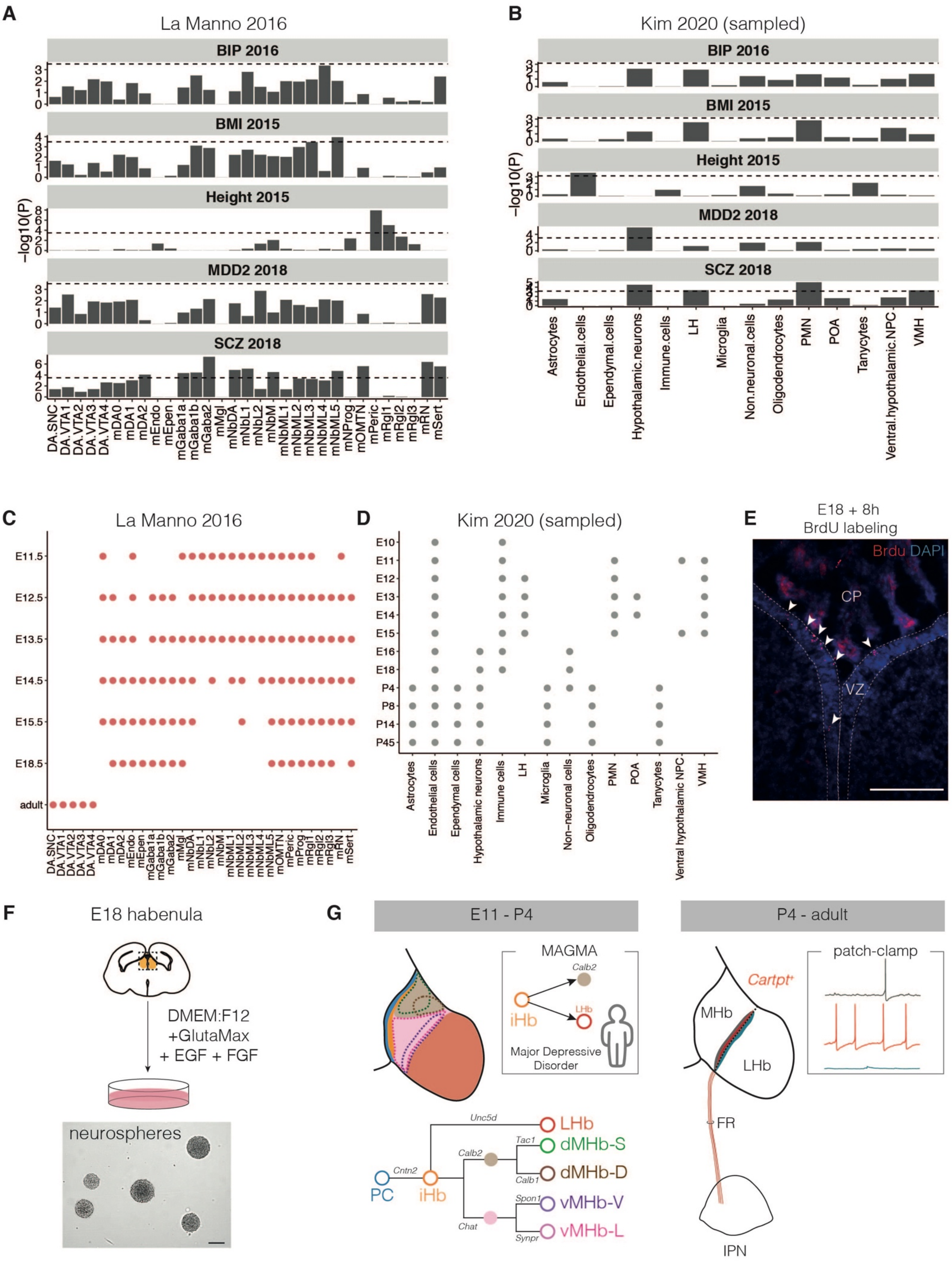

**Supplementary Figure 10. MAGMA analysis of developing midbrain and hypothalamus cell types. Related to Figure 8.**
**A, B,** Results of MAGMA analysis of midbrain dopamine cell types (La Manno et al., 2016) (A) and developing hypothalamus cell types (Kim et al., 2020). Bar plots resemble signal strength, if dashed line (*P* value after Bonferroni correction = 0.05 / (31*5)) (a) and = 0.05 / (14*5)) (b) is crossed, signal is considered significant.
**C, D,** Dot plots showing the age occupation per cell type per dataset.
**E,** E18 + 8 hours BrdU labeling. Arrows indicate $BrdU^+$ cells in the ventricular zone of the habenula.
**F,** Top, overview of methods for neurosphere culture of E18 habenula tissue. Bottom, brightfield image of neurospheres at DIV7. Scale bar = 100 μm.
BIP, Bipolar Disorder; BMI, Body-Mass Index; BrdU, Bromodeoxyuridine; CP, choroid plexus; DA, dopaminergic; DIV, days *in vitro*; E, embryonic day; EGF, epidermal growth factor; Endo, endothelial cells; Epend, ependymal; FGF, fibroblast growth factor; Gaba1-2, GABAergic neurons; LH, lateral hypothalamus; m, mouse; MDD, Major Depressive Disorder; Mgl, microglia; mNbL1-2, lateral neuroblasts; NbML1-5, mediolateral neuroblasts; NPC, neural precursor cell; OMTN, oculomotor and trochlear nucleus; PC, progenitor cells; Peric, pericytes; PMN, premammillary hypothalamus; Rgl1-3, radial glia-like cells; RN, red nucleus; SCZ, Schizophrenia; Sert, serotonergic; SNC, substantia nigra pars compacta; SVZ, subventricular zone; VMH, ventromedial hypothalamus; VTA, ventral tegmental area; VZ, ventricular zone.

**Supplementary Table S1.** Table provided as separate excel file.

**Supplementary Table S1. Top 99 differentially expressed genes per Louvain cluster of the developmental habenula dataset. Ranked based on statistical significance. Related to Figure 1.**

#### Supplementary Table S2.

| | scores | names | logfoldchanges | pvals | pvals_adj |
|---|---|---|---|---|---|
| 0 | 12,186506 | Hbb-bs | 1,7411668 | 3,67E-34 | 1,04E-29 |
| 1 | 11,354883 | Hba-a2 | 1,7982825 | 7,01E-30 | 9,90E-26 |
| 2 | 11,155998 | Hba-a1 | 1,6159922 | 6,69E-29 | 6,30E-25 |
| 3 | 9,099599 | Hbb-bt | 1,9299886 | 9,07E-20 | 6,40E-16 |
| 4 | 6,4834695 | Srgn | 5,391131 | 8,96E-11 | 5,06E-07 |
| 5 | 4,5760236 | H19 | 2,981257 | 4,74E-06 | 0,01911632 |
| 6 | 4,4195175 | Bsg | 0,5778652 | 9,89E-06 | 0,02793247 |
| 7 | 4,372524 | Haghl | 0,8625455 | 1,23E-05 | 0,0297452 |
| 8 | 4,36623 | Gm20521 | 0,45830095 | 1,26E-05 | 0,0297452 |
| 9 | 4,323432 | Serinc3 | 0,56837493 | 1,54E-05 | 0,03336757 |
| 10 | 4,23364 | Rpl30 | 0,57876664 | 2,30E-05 | 0,04637701 |
| 11 | 4,195038 | 2900060B14Rik | 1,2383692 | 2,73E-05 | 0,05051984 |
| 12 | 4,128743 | Rpl39 | 0,62325805 | 3,65E-05 | 0,05560944 |
| 13 | 4,122869 | Gm22710 | 1,2598829 | 3,74E-05 | 0,05560944 |
| 14 | 4,1060853 | Sars | 0,5512003 | 4,02E-05 | 0,05681576 |
| 15 | 4,093078 | Gm9844 | 0,44223803 | 4,26E-05 | 0,05723828 |
| 16 | 4,074197 | Blcap | 0,58791226 | 4,62E-05 | 0,05823958 |
| 17 | 4,067903 | Rpl36 | 0,5513228 | 4,74E-05 | 0,05823958 |
| 18 | 4,0146155 | Rps29 | 0,37846014 | 5,95E-05 | 0,06992936 |
| 19 | 3,988601 | Hdgfl3 | 0,45823935 | 6,65E-05 | 0,06992936 |
| 20 | 3,984405 | Lgals9 | 4,01902 | 6,76E-05 | 0,06992936 |
| 21 | 3,9835658 | Apoe | 1,2783098 | 6,79E-05 | 0,06992936 |
| 22 | 3,9625864 | Btbd11 | 0,7148567 | 7,41E-05 | 0,07152279 |
| 23 | 3,9567122 | Camta1 | 0,32800618 | 7,60E-05 | 0,07152279 |
| 24 | 3,9470618 | Ptprcap | 4,177963 | 7,91E-05 | 0,07206462 |
| 25 | 3,933635 | Dpysl4 | 0,60037607 | 8,37E-05 | 0,0723483 |
| 26 | 3,9311173 | B2m | 1,6470807 | 8,46E-05 | 0,0723483 |
| 27 | 3,8635638 | Meis1 | 2,5647986 | 0,00011174 | 0,09040185 |
| 28 | 3,8606267 | Def6 | 1,122947 | 0,0001131 | 0,09040185 |
| 29 | 3,850137 | Rbpms | 2,8470023 | 0,00011805 | 0,09040185 |
| 30 | 3,8492978 | Vps9d1 | 0,6558841 | 0,00011846 | 0,09040185 |
| 31 | 3,8119545 | Naca | 0,5253595 | 0,00013787 | 0,10244998 |
| 32 | 3,8027236 | Ica1 | 0,59512883 | 0,00014311 | 0,10294982 |
| 33 | 3,7863595 | Pabpn1 | 0,43464923 | 0,00015287 | 0,10294982 |
| 34 | 3,78594 | Rps27a | 0,35325825 | 0,00015313 | 0,10294982 |
| 35 | 3,7582471 | Mical3 | 0,6105946 | 0,00017111 | 0,11236209 |
| 36 | 3,7133512 | Retreg2 | 0,5063067 | 0,00020453 | 0,1312588 |
| 37 | 3,649574 | Rpl14 | 0,38592997 | 0,00026268 | 0,16124281 |
| 38 | 3,6420214 | Mta1 | 0,4944075 | 0,00027051 | 0,16251627 |
| 39 | 3,6071956 | Tlk1 | 0,5818195 | 0,00030952 | 0,17836809 |
| 40 | 3,5983844 | Rpl18 | 0,40659073 | 0,0003202 | 0,18077415 |
| 41 | 3,5899925 | Wdr7 | 0,47147036 | 0,00033069 | 0,18077415 |

| | | | | | |
|---|---|---|---|---|---|
| 42 | 3,5832791 | Atp1a3 | 0,53875345 | 0,00033931 | 0,18077415 |
| 43 | 3,5669153 | Acvr2b | 0,5331722 | 0,00036121 | 0,18633675 |
| 44 | 3,5362854 | Syp | 0,4891434 | 0,0004058 | 0,20102571 |
| 45 | 3,527474 | Ufc1 | 0,6702597 | 0,00041954 | 0,2042533 |
| 46 | 3,515306 | Sbf1 | 0,5594534 | 0,00043925 | 0,20656503 |
| 47 | 3,5127885 | Ttc1 | 0,73422366 | 0,00044343 | 0,20656503 |
| 48 | 3,51111 | Hmga2 | 1,7075582 | 0,00044624 | 0,20656503 |
| 49 | 3,498103 | Selenon | 0,67648906 | 0,00046858 | 0,20710992 |
| 50 | 3,4934874 | Ttc3 | 0,18562452 | 0,00047676 | 0,20710992 |
| 51 | 3,4934874 | Gnao1 | 0,26755193 | 0,00047676 | 0,20710992 |
| 52 | 3,472508 | Herpud1 | 0,79979 | 0,00051562 | 0,2196967 |
| 53 | 3,4695709 | Pnck | 0,48760206 | 0,00052129 | 0,2196967 |
| 54 | 3,4523678 | Mef2c | 1,1030918 | 0,00055569 | 0,23075015 |
| 55 | 3,4422977 | Znhit1 | 0,79526144 | 0,0005768 | 0,23604296 |
| 56 | 3,4238358 | Mecom | 1,3626434 | 0,00061744 | 0,24555813 |
| 57 | 3,406213 | Grk2 | 0,54361004 | 0,00065871 | 0,24991157 |
| 58 | 3,4041152 | Camk2d | 0,5001506 | 0,00066379 | 0,24991157 |
| 59 | 3,3919473 | Gfpt1 | 0,50539696 | 0,00069398 | 0,2578403 |
| 60 | 3,3843946 | Flywch2 | 0,74909633 | 0,00071335 | 0,26159712 |
| 61 | 3,376842 | Ncam1 | 0,15965791 | 0,00073323 | 0,26543922 |
| 62 | 3,3663523 | Sytl2 | 0,8867348 | 0,00076169 | 0,26553014 |
| 63 | 3,357541 | Ubqln2 | 0,3953966 | 0,00078639 | 0,26957141 |
| 64 | 3,355443 | 4933434E20Rik | 0,4838048 | 0,00079238 | 0,26957141 |
| 65 | 3,3499885 | Dennd5b | 0,52240413 | 0,00080815 | 0,27166329 |
| 66 | 3,3374007 | Sec61a2 | 0,49170923 | 0,00084566 | 0,28092797 |
| 67 | 3,331107 | Tenm1 | 0,6455876 | 0,00086501 | 0,28401616 |
| 68 | 3,3080297 | Timm29 | 0,5502638 | 0,00093955 | 0,3014777 |
| 69 | 3,304673 | Meg3 | 0,5263806 | 0,00095087 | 0,30168315 |
| 70 | 3,2950225 | Slc35f1 | 0,61093235 | 0,00098414 | 0,30537489 |
| 71 | 3,2769802 | Pde4a | 0,65179104 | 0,00104924 | 0,31706293 |
| 72 | 3,2753017 | Dym | 0,72647583 | 0,00105549 | 0,31706293 |
| 73 | 3,2753017 | 2410089E03Rik | 0,50838804 | 0,00105549 | 0,31706293 |
| 74 | 3,2610357 | Lmo2 | 1,2882603 | 0,00111006 | 0,32844597 |
| 75 | 3,2593575 | Atp13a2 | 0,3848049 | 0,00111665 | 0,32844597 |
| 76 | 3,2358606 | Cnot6l | 0,51098955 | 0,00121277 | 0,35304001 |
| 77 | 3,2249513 | Tcea2 | 0,7960977 | 0,00125994 | 0,36165366 |
| 78 | 3,2203357 | Stxbp2 | 0,789851 | 0,00128041 | 0,36165366 |
| 79 | 3,2194967 | Gm26694 | 0,5672217 | 0,00128416 | 0,36165366 |
| 80 | 3,2111049 | Csf2ra | 0,48149088 | 0,00132226 | 0,36604475 |
| 81 | 3,1972585 | Cbfa2t3 | 0,774492 | 0,00138741 | 0,37669394 |
| 82 | 3,1871884 | Gpx4 | 0,6680555 | 0,00143663 | 0,38634451 |
| 83 | 3,1678872 | Rpl36a | 0,34886134 | 0,00153551 | 0,4000841 |
| 84 | 3,1670482 | Samd12 | 0,47198188 | 0,00153995 | 0,4000841 |
| 85 | 3,166209 | Nrsn1 | 0,7934082 | 0,0015444 | 0,4000841 |
| 86 | 3,1569781 | Txnip | 0,98294 | 0,00159413 | 0,40364507 |
| 87 | 3,1557193 | Lig1 | 0,66177577 | 0,00160103 | 0,40364507 |
| 88 | 3,1494255 | Nnat | 0,27920407 | 0,00163592 | 0,40398318 |
| 89 | 3,147747 | Rnf40 | 0,7242215 | 0,00164534 | 0,40398318 |
| 90 | 3,1452296 | Gab2 | 0,6371483 | 0,00165957 | 0,40398318 |
| 91 | 3,1443903 | Pygo1 | 0,6560147 | 0,00166433 | 0,40398318 |

| | | | | | |
|---|---|---|---|---|---|
| 92 | 3,142712 | Trpm3 | 0,4948784 | 0,0016739 | 0,40398318 |
| 93 | 3,1271873 | Rcan2 | 0,5875652 | 0,00176487 | 0,41886004 |
| 94 | 3,1259286 | Fmnl1 | 0,99493825 | 0,00177245 | 0,41886004 |
| 95 | 3,1217327 | B3gat1 | 0,41033208 | 0,0017979 | 0,41956475 |
| 96 | 3,1150193 | Lsamp | 0,3622721 | 0,00183933 | 0,4257142 |
| 97 | 3,1024315 | Zbtb8b | 1,2252358 | 0,00191938 | 0,4370766 |
| 98 | 3,0894244 | Usp29 | 0,4422449 | 0,00200545 | 0,45286577 |
| 99 | 3,084809 | Fam110b | 0,72920257 | 0,00203683 | 0,45286577 |

**Supplementary Table S2. Top 99 Differentially expressed genes in the left hemisphere of the E18 WT habenula compared to the right hemisphere. Ranked based on statistical significance. Related to Figure 5.**

**Supplementary Table S3**

| | scores | names | logfoldchanges | pvals | pvals_adj |
|---|---|---|---|---|---|
| 0 | 6,233395 | Gm12504 | 0,98770463 | 4,56E-10 | 2,58E-06 |
| 1 | 5,0522556 | Ybx1 | 0,3115542 | 4,37E-07 | 0,00176127 |
| 2 | 4,6700115 | Msi1 | 0,5743762 | 3,01E-06 | 0,01063063 |
| 3 | 4,5437155 | Selenok | 0,45255154 | 5,53E-06 | 0,01734105 |
| 4 | 4,433364 | Marcksl1 | 0,21028297 | 9,28E-06 | 0,02619662 |
| 5 | 4,2025905 | Basp1 | 0,29413393 | 2,64E-05 | 0,06059025 |
| 6 | 4,190003 | Nedd4 | 0,20336275 | 2,79E-05 | 0,06059025 |
| 7 | 4,088463 | Uba1 | 0,20670268 | 4,34E-05 | 0,08758335 |
| 8 | 4,0171328 | Tmsb4x | 0,23588839 | 5,89E-05 | 0,10412914 |
| 9 | 4,004545 | Ptn | 0,8835995 | 6,21E-05 | 0,10412914 |
| 10 | 3,8761513 | Hnrnpa2b1 | 0,19123094 | 0,00010612 | 0,13028511 |
| 11 | 3,8648226 | Ppp2r1a | 0,24465673 | 0,00011117 | 0,13079614 |
| 12 | 3,8249617 | Gpm6b | 0,41416138 | 0,00013079 | 0,14759721 |
| 13 | 3,7884576 | Hpcal1 | 0,4405889 | 0,00015159 | 0,14759721 |
| 14 | 3,7603452 | Rbbp4 | 0,37373602 | 0,00016968 | 0,15970759 |
| 15 | 3,7091556 | Ndufc2 | 0,3337643 | 0,00020795 | 0,1777466 |
| 16 | 3,6890154 | Hsf3 | 0,6728618 | 0,00022512 | 0,1777466 |
| 17 | 3,687337 | Cltc | 0,31694028 | 0,00022661 | 0,1777466 |
| 18 | 3,5908318 | Pfdn2 | 0,45190385 | 0,00032962 | 0,20683564 |
| 19 | 3,5362854 | Hmgcs1 | 0,4056583 | 0,0004058 | 0,22818233 |
| 20 | 3,5270543 | Eid1 | 0,28830445 | 0,00042021 | 0,22818233 |
| 21 | 3,4414585 | Elob | 0,29221568 | 0,00057859 | 0,30254752 |
| 22 | 3,4120874 | Snap47 | 0,26442719 | 0,00064467 | 0,31380236 |
| 23 | 3,3743246 | Bex3 | 0,23055224 | 0,00073997 | 0,34824263 |
| 24 | 3,3684502 | Tmed2 | 0,564623 | 0,00075592 | 0,34991678 |
| 25 | 3,341177 | Otx1 | 0,6952229 | 0,00083424 | 0,37391169 |
| 26 | 3,303414 | Abhd2 | 0,7071336 | 0,00095515 | 0,41185516 |
| 27 | 3,2967007 | Gid8 | 0,82289803 | 0,00097828 | 0,41185516 |
| 28 | 3,295442 | Rpl15 | 0,3170665 | 0,00098267 | 0,41185516 |
| 29 | 3,2585182 | Elp2 | 0,3338591 | 0,00111996 | 0,43922522 |
| 30 | 3,2417347 | Ndufb6 | 0,42520747 | 0,00118805 | 0,4525879 |
| 31 | 3,238378 | Eif4g2 | 0,17548518 | 0,00120211 | 0,4525879 |
| 32 | 3,1825728 | Kdm1a | 0,40976986 | 0,00145973 | 0,49000998 |
| 33 | 3,1737616 | Ptms | 0,21106745 | 0,00150477 | 0,49000998 |
| 34 | 3,1594956 | Wsb1 | 0,32339036 | 0,00158043 | 0,49000998 |
| 35 | 3,1586564 | Hnrnpc | 0,21299954 | 0,00158498 | 0,49000998 |
| 36 | 3,15488 | E130308A19Rik | 0,33480677 | 0,00160564 | 0,49000998 |
| 37 | 3,1527822 | Hdac2 | 0,3450788 | 0,00161722 | 0,49000998 |
| 38 | 3,1527822 | Rtraf | 0,3226305 | 0,00161722 | 0,49000998 |
| 39 | 3,1511037 | Copb1 | 0,49551913 | 0,00162655 | 0,49000998 |
| 40 | 3,1460688 | Ube2e3 | 0,26561892 | 0,00165481 | 0,49186208 |
| 41 | 3,1292853 | Rnf5 | 0,33047223 | 0,00175232 | 0,51010609 |
| 42 | 3,1175368 | Wls | 0,47650677 | 0,00182369 | 0,52015744 |
| 43 | 3,111243 | Gnb2 | 0,34490564 | 0,00186302 | 0,52605976 |
| 44 | 3,0982358 | Phf21b | 0,7115558 | 0,00194676 | 0,539693 |
| 45 | 3,0831306 | Arl8a | 0,32709885 | 0,00204835 | 0,55614737 |
| 46 | 3,0759976 | Mtss1l | 0,31004372 | 0,002098 | 0,55756325 |
| 47 | 3,061312 | Ehbp1 | 0,31094125 | 0,00220369 | 0,56611613 |

| | | | | | |
|---|---|---|---|---|---|
| 48 | 3,0537596 | Mmgt1 | 0,4477147 | 0,00225993 | 0,56976519 |
| 49 | 3,0336192 | Zic1 | 0,17485924 | 0,00241639 | 0,58820395 |
| 50 | 3,0302625 | Cmpk1 | 0,30803594 | 0,00244341 | 0,58969773 |
| 51 | 3,02271 | 2610307P16Rik | 0,5671429 | 0,00250522 | 0,59610385 |
| 52 | 3,0218709 | Eif1 | 0,14399911 | 0,00251218 | 0,59610385 |
| 53 | 3,0185142 | Cnn3 | 0,55908674 | 0,00254018 | 0,59772456 |
| 54 | 3,0038285 | Gm37121 | 0,50354564 | 0,00266606 | 0,60880543 |
| 55 | 2,9815903 | Hsp90ab1 | 0,18157005 | 0,00286755 | 0,60880543 |
| 56 | 2,9815903 | Zcrb1 | 0,26832294 | 0,00286755 | 0,60880543 |
| 57 | 2,9622893 | Pcca | 0,40558165 | 0,00305361 | 0,62456176 |
| 58 | 2,9530585 | Prmt2 | 0,23038632 | 0,00314642 | 0,62808738 |
| 59 | 2,9522192 | Eif2b5 | 0,6442045 | 0,00315499 | 0,62808738 |
| 60 | 2,9429884 | Snrpn | 0,15202737 | 0,00325061 | 0,63038701 |
| 61 | 2,9425688 | Tmem176b | 0,35675716 | 0,00325502 | 0,63038701 |
| 62 | 2,9421492 | Cbln4 | 0,7849582 | 0,00325943 | 0,63038701 |
| 63 | 2,9354358 | Zic3 | 0,4362101 | 0,0033308 | 0,63980742 |
| 64 | 2,9287224 | Rdh5 | 0,39990973 | 0,00340358 | 0,64937151 |
| 65 | 2,9169738 | Csnk1d | 0,25589094 | 0,00353445 | 0,66534917 |
| 66 | 2,9043863 | Tbl1xr1 | 0,48904407 | 0,00367974 | 0,6791163 |
| 67 | 2,8917985 | Nrn1 | 0,20024258 | 0,00383043 | 0,68891699 |
| 68 | 2,8884418 | Dynlrb1 | 0,19622833 | 0,00387156 | 0,69190578 |
| 69 | 2,855714 | Cacybp | 0,19074756 | 0,00429402 | 0,74546866 |
| 70 | 2,850679 | Mlf2 | 0,17444992 | 0,0043626 | 0,74546866 |
| 71 | 2,8490007 | Tagln3 | 0,20607504 | 0,00438568 | 0,74546866 |
| 72 | 2,8490007 | Nfib | 0,56319356 | 0,00438568 | 0,74546866 |
| 73 | 2,8481615 | Tubb5 | 0,15699512 | 0,00439726 | 0,74546866 |
| 74 | 2,8301191 | Atp5g3 | 0,1711499 | 0,00465307 | 0,74564926 |
| 75 | 2,8221471 | Uchl1 | 0,10990852 | 0,00477033 | 0,74833228 |
| 76 | 2,8162727 | Smarcd1 | 0,33970574 | 0,00485844 | 0,75794336 |
| 77 | 2,807042 | Kirrel3 | 0,5811439 | 0,00499987 | 0,76729047 |
| 78 | 2,807042 | Ralgps1 | 0,36962482 | 0,00499987 | 0,76729047 |
| 79 | 2,7919366 | Otx2 | 0,5083793 | 0,00523936 | 0,78267479 |
| 80 | 2,7860625 | Arf4 | 0,24800487 | 0,00533526 | 0,78267479 |
| 81 | 2,7860625 | Irx1 | 0,28055814 | 0,00533526 | 0,78267479 |
| 82 | 2,7827058 | Glrx3 | 0,42864656 | 0,00539077 | 0,78267479 |
| 83 | 2,7801883 | Disp3 | 0,79870284 | 0,00543274 | 0,78267479 |
| 84 | 2,766342 | Polr1d | 0,37382308 | 0,00566891 | 0,80438659 |
| 85 | 2,7617264 | Mir678 | 0,28291503 | 0,00574966 | 0,81176628 |
| 86 | 2,7474604 | Sh3rf1 | 0,6252333 | 0,00600588 | 0,83954413 |
| 87 | 2,7373903 | Klhl9 | 0,30355453 | 0,00619288 | 0,85561238 |
| 88 | 2,7331944 | Kctd5 | 0,6362139 | 0,00627233 | 0,85561238 |
| 89 | 2,7331944 | Dynlt1f | 0,65282816 | 0,00627233 | 0,85561238 |
| 90 | 2,726481 | Arl2bp | 0,23238197 | 0,00640136 | 0,86901582 |
| 91 | 2,7189283 | Mrps28 | 0,22742282 | 0,00654938 | 0,88064174 |
| 92 | 2,7046626 | Marcks | 0,11582422 | 0,00683738 | 0,90641863 |
| 93 | 2,6945925 | Olfm2 | 0,2549159 | 0,00704748 | 0,91096509 |
| 94 | 2,6937532 | Fscn1 | 0,13701811 | 0,00706525 | 0,91096509 |
| 95 | 2,6937532 | Arf2 | 0,28051168 | 0,00706525 | 0,91096509 |
| 96 | 2,6874595 | Abraxas2 | 0,5607873 | 0,00719979 | 0,92409253 |
| 97 | 2,6752913 | Sulf2 | 0,45955417 | 0,00746643 | 0,94644574 |

| | | | | | |
|---|---|---|---|---|---|
| 98 | 2,6719346 | Hacd3 | 0,2938641 | 0,00754153 | 0,94644574 |
| 99 | 2,6648016 | Zc4h2 | 0,4804486 | 0,00770337 | 0,95105557 |

**Supplementary Table S3. Top 99 Differentially expressed genes in the right hemisphere of the E18 WT habenula compared to the left hemisphere. Ranked based on statistical significance. Related to Figure 5.**

#### Supplementary Table S4

| Genotyping | | | | |
|---|---|---|---|---|
| **Locus** | **forward primer** | **reverse primer** | **size (bp)** | |
| *Brn3a-wt* | ACA AGC CGG AGC TCT TCA ACG G | CAG GAT AAC GGA CAG TCT AAA TGA | 330 | |
| *Brn3a-tLacZ* | TATCCGAACCATCCGCTGTG | TTAGCGAAACCGCCAAGACT | 479 | |
| *Cart-IRES2-cre-WT* | CCA TCG GTT GTG TTG TTG AC | TCG AGG CAT TCT CCT TCA CAC | 315 | |
| *Cart-IRES2-cre-mutant* | CCA TCG GTT GTG TTG TTG AC | ACA CCG GCC TTA TTC CAA G | 200 | |
| *Cartpt-cre-ERT2-wt* | GCTGCCTACAGACGGCTGAC | GGAGCTCTCCATGGTTCTGG | 172 | |
| *Cartpt-cre-ERT2-mutant* | GCTGCCTACAGACGGCTGAC | ACATCTTCAGGTTCTGCG | 430 | |
| *Td-Tomato WT* | AAGGGAGCTGCAGTGGAGTA | CCGAAAATCTGTGGGAAGTC | 279 | |
| *Td-Tomato mutant* | GGCATTAAAGCAGCGTATCC | CTGTTCCTGTACGGCATGG | 196 | |
| ***In situ* hybridization probes** | | | | |
| **probe** | **primer_FW** | **primer_RV** | **size (bp)** | **transcript** |
| Tac1 | CCCCTGAACGCACTATCTATTC | CAGGAAACATGCTGCTAGGATA | 428 | NM_009311.3 |
| Kcng4 | AAACAAGCGACAGGTGAGCC | AGGTTGGTTTCCTCGATGCC | 517 | NM_025734.2 |
| Etv1 | GTGCCTCTGTCTCACTTTGATG | CTACTGGCCTGTGACTCAGTTG | 804 | NM_007960.5 |
| Cntn2 | ATCTAGTGTCTGGGCAGGGTA A | TTTATAGGCAGCTTCACCCCTA | 437 | NM_177129.5 |
| Nrgn | AGAGGAGAGAGGCTGGTTCTG | GCAAAAGGAGACCACGTTAGAC | 616 | NM_022029.2 |
| Id3 | AGGTGTCTCTTTTCCTCCCTCT | TCTCCAAGGAAACCAGAAGAAC | 783 | NM_008321.2 |
| Dbi | TCGGCATCCGTATCACCT | GAGGTTAACGCTGGCCCT | 399 | NM_001037999.2 |
| Nwd2 | ATGTGCTGGATTTACACAGTG G | CCAGAAATTAGCAGCTCATCA | 691 | NM_177006.3 |

**Supplementary Table S4. Overview of oligonucleotides used for genotyping and creation of the in situ hybridization probes. Related to Figure 1, 5 and 6.**
